# Prototype-based continual cell-type annotation reveals cellular state transitions in expanding single-cell atlases

**DOI:** 10.64898/2026.03.05.709973

**Authors:** Shuang Ge, Qiming He, Yiming Ren, Yaxin Xu, Mingqing Wang, Zhiwei Nie, Huan Xu, Qiang Cheng, Shuqing Sun, Zhixiang Ren

## Abstract

Large-scale single-cell atlases provide an increasingly comprehensive view of cellular diversity, but their continued expansion across studies poses a fundamental challenge: preserving consistent cell identities while capturing biological variation in cellular states. Most existing annotation frameworks are built around static references, making it difficult to incorporate newly generated datasets into established cellular representations without retraining on historical data. When updated sequentially, these methods are further constrained by catastrophic forgetting and batch-specific biases, limiting scalability and the continuity of knowledge integration. Here we introduce scEvolver, a continual learning framework for single-cell annotation that incrementally accumulates knowledge through memory-guided refinement of cell-type prototypes without revisiting historical data. Across sequencing platforms, tissue contexts and molecular modalities, scEvolver supports robust annotation and external query mapping with substantially fewer labelled reference cells. By preserving consistent cell-type semantics across datasets while capturing biologically meaningful within-class heterogeneity, scEvolver enables the identification of epithelial cell-state transitions in inflammatory gut disease. External mapping to the healthy Human Lung Cell Atlas further reveals shared cell-state deviations across multiple diseases, including an FCGR3A^+^ inflammatory monocyte programme in sarcoidosis, chronic obstructive pulmonary disease and idiopathic pulmonary fibrosis, highlighting scEvolver’s potential to characterize context-specific cellular dynamics in complex disease settings.

## 1 Introduction

Single-cell sequencing has improved our understanding of cellular heterogeneity and enabled the construction of large-scale single-cell atlases for biological discovery Jovic et al. [2022], Wen and Tang [2022], Ge et al. [2025]. Within this context, cell-type annotation serves as a core task that assigns expression profiles to specific cell identities, providing critical insights into cellular states and functions Vieth et al. [2019], Luecken and Theis [2019]. However, traditional annotation approaches rely on predefined markers and expert expertise, making them time-consuming and difficult to scale Cheng et al. [2023]. In addition, heterogeneity in experimental protocols and tissue origins introduces systematic biases, hindering consistent annotation across datasets Yu et al. [2024], Zhou et al. [2025], Baysoy et al. [2023], Sun et al. [2025]. As single-cell datasets grow in scale, severe class imbalance and limited reference annotations further increase the difficulty of accurate annotation Zhao et al. [2020]. In addition, privacy constraints and computational costs make retraining models from scratch increasingly impractical, challenging effective knowledge integration and generalization Walker et al. [2024], Chen et al. [2023].

To overcome the limitations of manual annotation, researchers have developed a wide range of automated tools, ranging from traditional machine learning classifiers to advanced deep learning models Pasquini et al. [2021], Ma and Pellegrini [2020], Johnson et al. [2019], Cao et al. [2020], Kimmel and Kelley [2020]. More recently, large-scale foundation models have further improved generalization by learning complex biological patterns from millions of cells Yang et al. [2022], Cui et al. [2024]. Despite these advances, most existing methods are still designed for static datasets and struggle to accommodate the continual arrival of new data. When applied to sequential datasets, these models often suffer from catastrophic forgetting, whereby previously learned knowledge is lost as new information is integrated Ke et al. [2023]. Consequently, incorporating new datasets typically requires retraining models on the full reference set, which becomes increasingly costly and time-consuming as atlases grow Zhou et al. [2024], Zheng et al. [2025]. This requirement is further complicated by privacy regulations and data governance policies that restrict access to historical datasets. In addition to these limitations, existing models are sensitive to class imbalance, leading to biased predictions that favor abundant cell types over rare populations Khan et al. [2023]. Moreover, when reference annotations are relatively scarce, current frameworks often fail to learn sufficiently robust features, limiting their ability to generalize to novel biological contexts Cheng et al. [2023], Boiarsky et al. [2023].

To address these limitations, we leverage the dynamic evolution of cell type representations as class prototypes in a shared embedding space to incrementally integrate biological knowledge across expanding single-cell atlases. We introduce scEvolver, a prototype-based continual learning framework for single-cell annotation that adaptively updates cellular representations without revisiting past data. scEvolver integrates several key components to support robust annotation and analysis, including continual reference atlas construction, accurate query mapping into a harmonized latent space, reliable detection of outlier or novel cell populations, and the identification of prototype-correlated gene signatures that capture cellular heterogeneity and state transitions (Fig. 1). We evaluate scEvolver across a range of real-world scenarios, demonstrating unified cell representations across sequencing technologies (Fig. 2), robust cross-tissue generalization (Fig. 3), and accurate alignment of multimodal representations while effectively mitigating catastrophic forgetting and enabling continual knowledge accumulation (Fig. 4). Notably, scEvolver maintains high annotation accuracy even under extreme label scarcity (Fig. 5). Finally, we show that scEvolver captures biologically variation by linking gene expression to distances from class prototypes, supported by cell-type–specific markers and pathway enrichment analysis. Application of scEvolver to inflammatory gut disease data validates its ability to capture disease-associated metaplastic epithelial transitions, including the emergence of surface foveolar-like (SF-like) epithelial cells (Fig. 6). Reference mapping to healthy prototypes derived from the Human Lung Cell Atlas further shows that scEvolver can characterize shared disease-associated cell-state deviations across lung diseases and identify annotation-discordant populations in a prototype-guided manner (Fig. 7). These findings highlight the ability of scEvolver to capture subtle, biologically relevant cell-state changes in complex disease contexts, demonstrating its value for studying dynamic tissue remodeling and disease progression.

**Fig. 1.**
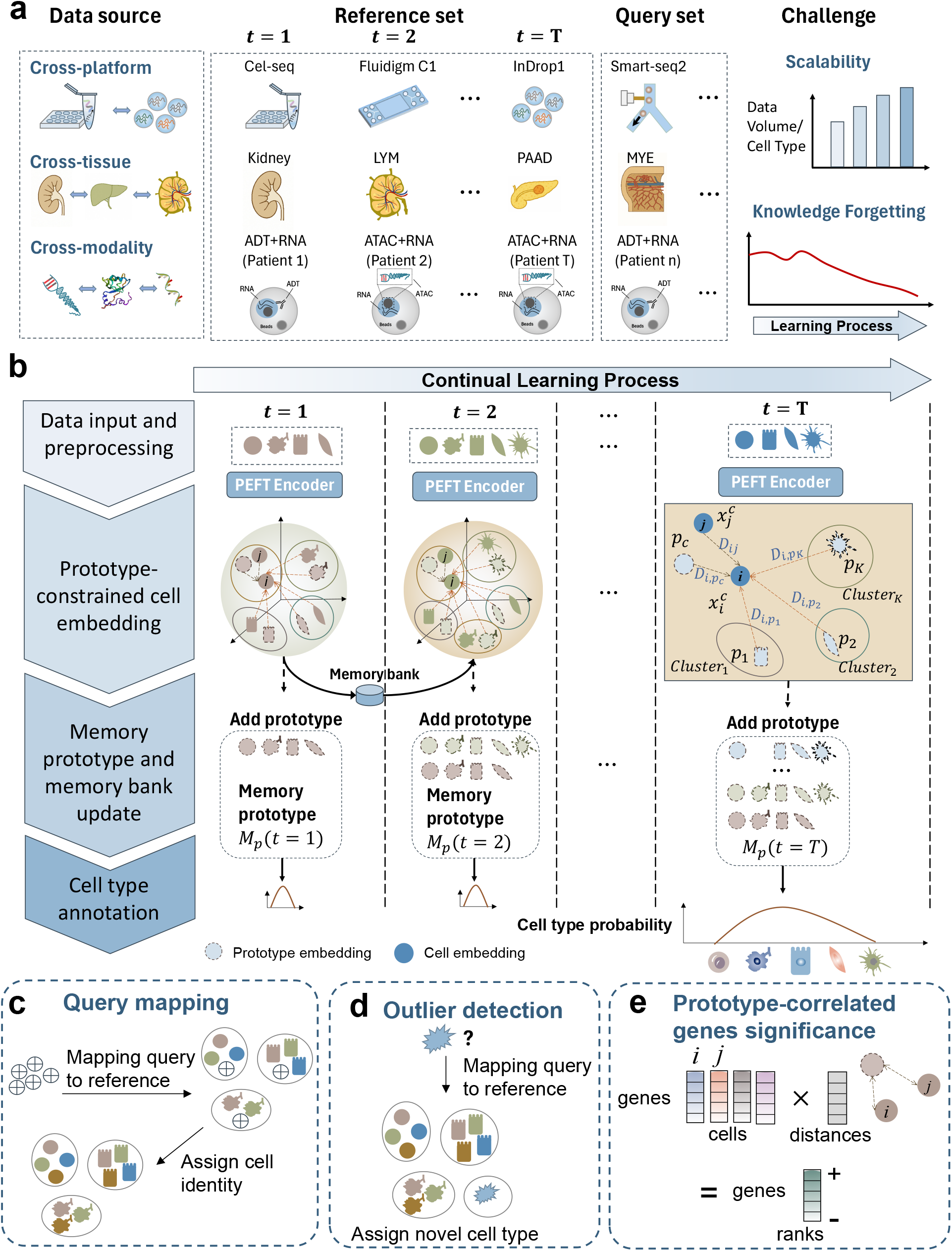
Overview of scEvolver. **a**, In real-world cell annotation scenarios, single-cell datasets are generated sequentially and exhibit substantial heterogeneity. Datasets may originate from different sequencing platforms (for example, CEL-Seq, Fluidigm C1 and 10x Genomics), span multiple tissues, or encompass diverse modalities. Sequential data acquisition and batch-wise distributional shifts introduce batch effects, while sequential learning settings are prone to catastrophic forgetting. **b**, The incremental annotated data comprising gene expression profiles and cell-type labels is preprocessed and feed into the model (step 1). Gene expression profiles are encoded into latent embeddings using a PEFT-based encoder. These embeddings are constrained by class prototypes in the latent space to reduce technical variation across datasets while preserving biologically relevant within-class heterogeneity, such that cells of the same type cluster around their corresponding prototypes (step 2). The model maintains a memory bank and updates memory prototypes to retain knowledge from previously learned data (step 3). Finally, we use a classifier to output the annotation results (step 4). **c**, ScEvolver supports query mapping under both online (including rare cell types) and few-shot online settings. **d, e**, scEvolver incorporates a similarity score between cells and their corresponding prototypes, enabling detection of unseen cell types and identification of prototype-correlated genes with either positive or negative associations.

**Fig. 2.**
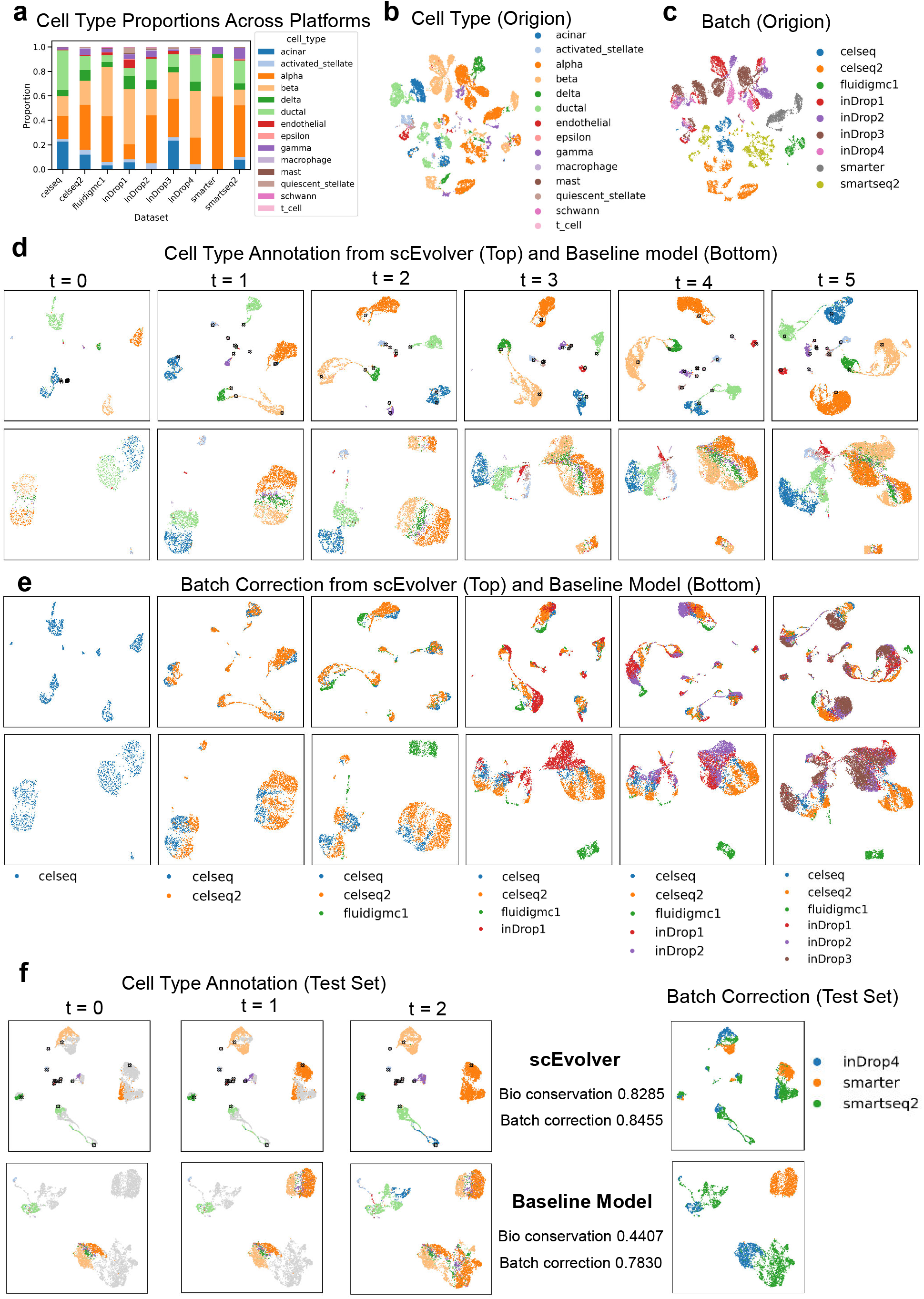
scEvolver harmonizes representations across sequencing platforms. **a**, Number of annotated cell types identified across different experimental platforms, showing variation in cell-type composition across batches. **b, c**, UMAP visualization of the original gene expression profiles with cell-type annotations (**b**) and batch labels (**c**), showing inconsistent alignment of the same cell types across platforms. **d**, UMAP visualization of cell embeddings during continual learning as new batches are incrementally incorporated. Cells of the same type from different sequencing platforms progressively align in the latent space, indicating effective integration across batches. Black crosses denote class prototypes, which evolve with incoming data while remaining well aligned with cell clusters, reflecting stable and semantically consistent class representations. **e**, UMAP visualization of batch mixing at incremental stages of continual learning. The baseline model exhibits clear batch separation, whereas scEvolver shows improved batch integration. **f**, Cell-type annotation results on held-out test datasets, evaluated after continual reference annotation. scEvolver reduces batch effects and yields clearer classification boundaries compared with the baseline model and other methods (supplementary Fig. S1 and S2; supplementary Table S2).

**Fig. 3.**
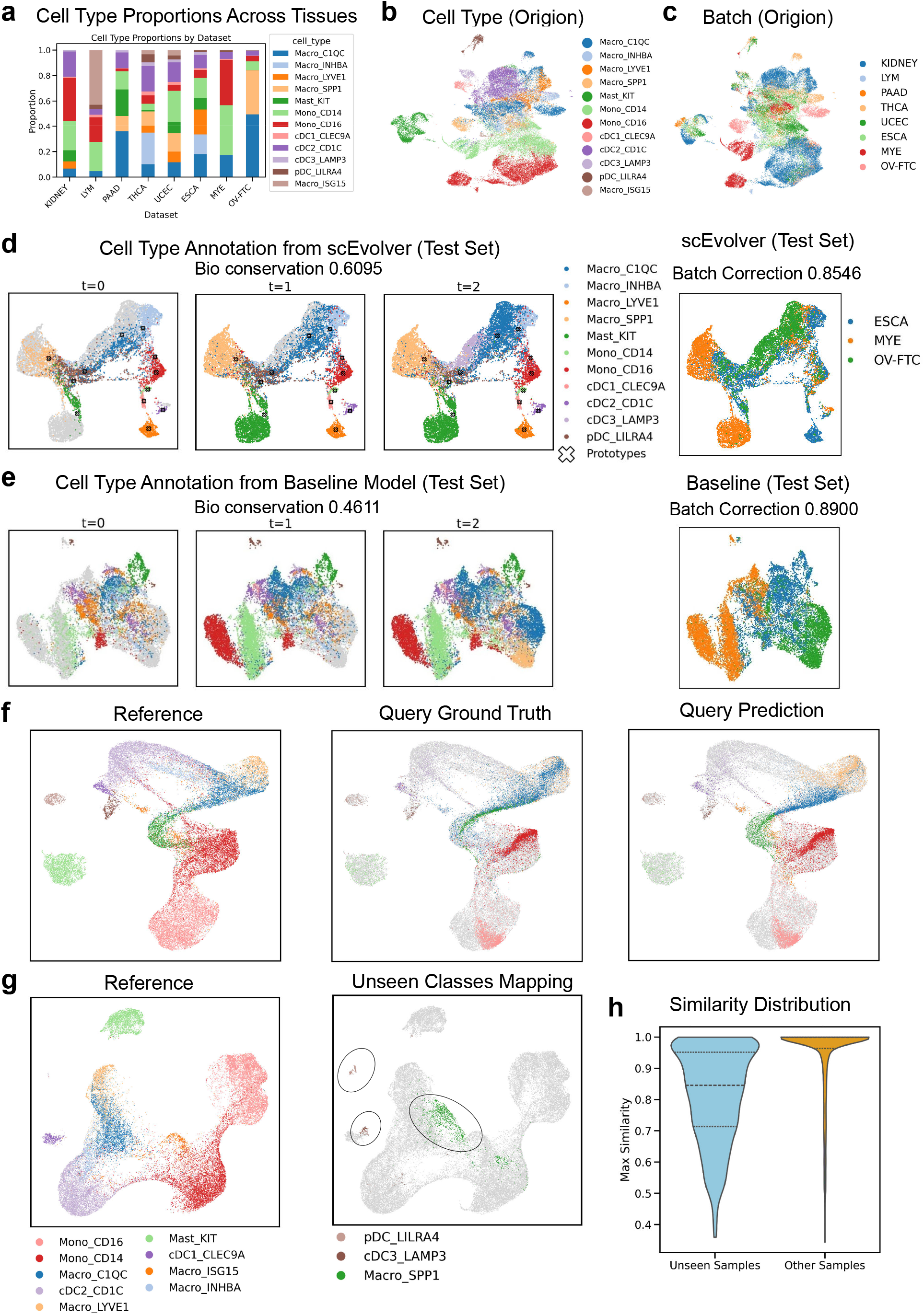
scEvolver enables generalizable query mapping and outlier detection. **a**, Number of annotated cell types identified across different tissues, illustrating tissue-specific differences in cell-type composition. **b, c**, UMAP visualization of the original gene expression profiles with cell-type annotations (**b**) and batch labels (**c**), showing inconsistent alignment of the same cell types across tissues. **d, e**, Cell-type annotation results on held-out test samples evaluated after successive stages of continual learning. scEvolver (**d**) shows improved cell-type organization and reduced batch-driven mixing compared with alternative annotation methods (**e**; Supplementary Fig. S1 and Fig. S2). **f**, UMAP visualization of reference data, query cells with ground-truth labels, and query cells with predicted annotations in the learned latent space, demonstrating accurate query mapping by scEvolver. **g**, UMAP visualization of the reference data after excluding the pDC_LILRA4, cDC3_LAMP3, and Macro_SPP1 cell types, along with the projection of these unseen cell types into the reference space. **h**, Distribution of maximum prototype similarity scores for known (Other Samples) and unseen cell types (Unseen Samples). Approximately 75% of unseen cells had similarity scores below 0.95, while more than 75% of known cells exceeded this threshold, demonstrating that unseen cell types remain distant from all existing class prototypes in the latent space. The three lines in the box plot represent the 25th, 50th, and 75th percentiles, respectively.

**Fig. 4.**
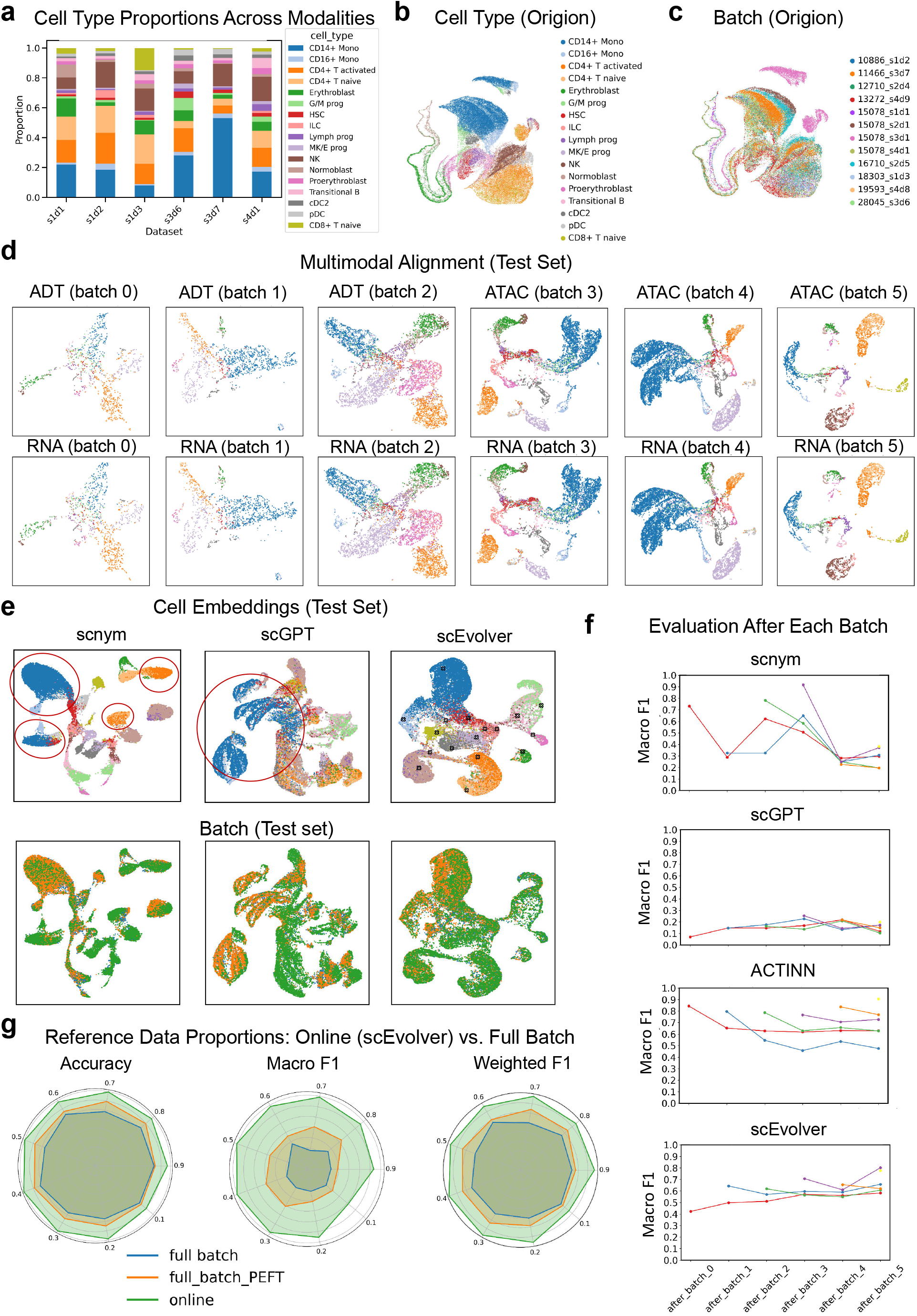
Continual cross-modal annotation with scEvolver mitigates catastrophic forgetting while enabling knowledge accumulation. **a**, Number of cell types across multimodal samples, each comprising paired RNA–ADT and RNA–ATAC measurements. **b, c**, UMAP visualization of the original gene expression profiles with cell-type annotations (**b**) and batch labels (**c**), showing inconsistent alignment of the same cell types across modalities. **d**, Alignment of multimodal representations in the latent space across successive batches (batches 0–5). **e**, Comparison of latent-space representations on the held-out test set. scNym and scGPT show separation among cells of the same type (highlighted by red circles), whereas scEvolver produces more coherent and continuous clustering. **f**, Forgetting curves showing model accuracy on previous datasets during continual learning. The x-axis represents the stages of data accumulation during continual learning. Each point on the curves represents the model’s test macro-F1 score on the current batch after completing training on the nth batch. The first point on the red curve, located at *after_batch_0*, represents the forgetting curve for the 0th batch. The blue curve represents the forgetting curve for the 1st batch, and so on. For example, the second point on the red curve represents the test score on data from “batch 0” after training on “batch 1”. **g**, Annotation performance across different proportions of labeled reference data (10–90%). *Full batch* indicates a baseline model trained jointly on all datasets, *Full_batch_PEFT* denotes PEFT fine-tuning on the complete dataset, and *Online* corresponds to the scEvolver continual learning approach. scEvolver achieves consistently strong performance across different proportions of labeled reference data, particularly in Macro F1 scores, highlighting its robustness to class imbalance.

**Fig. 5.**
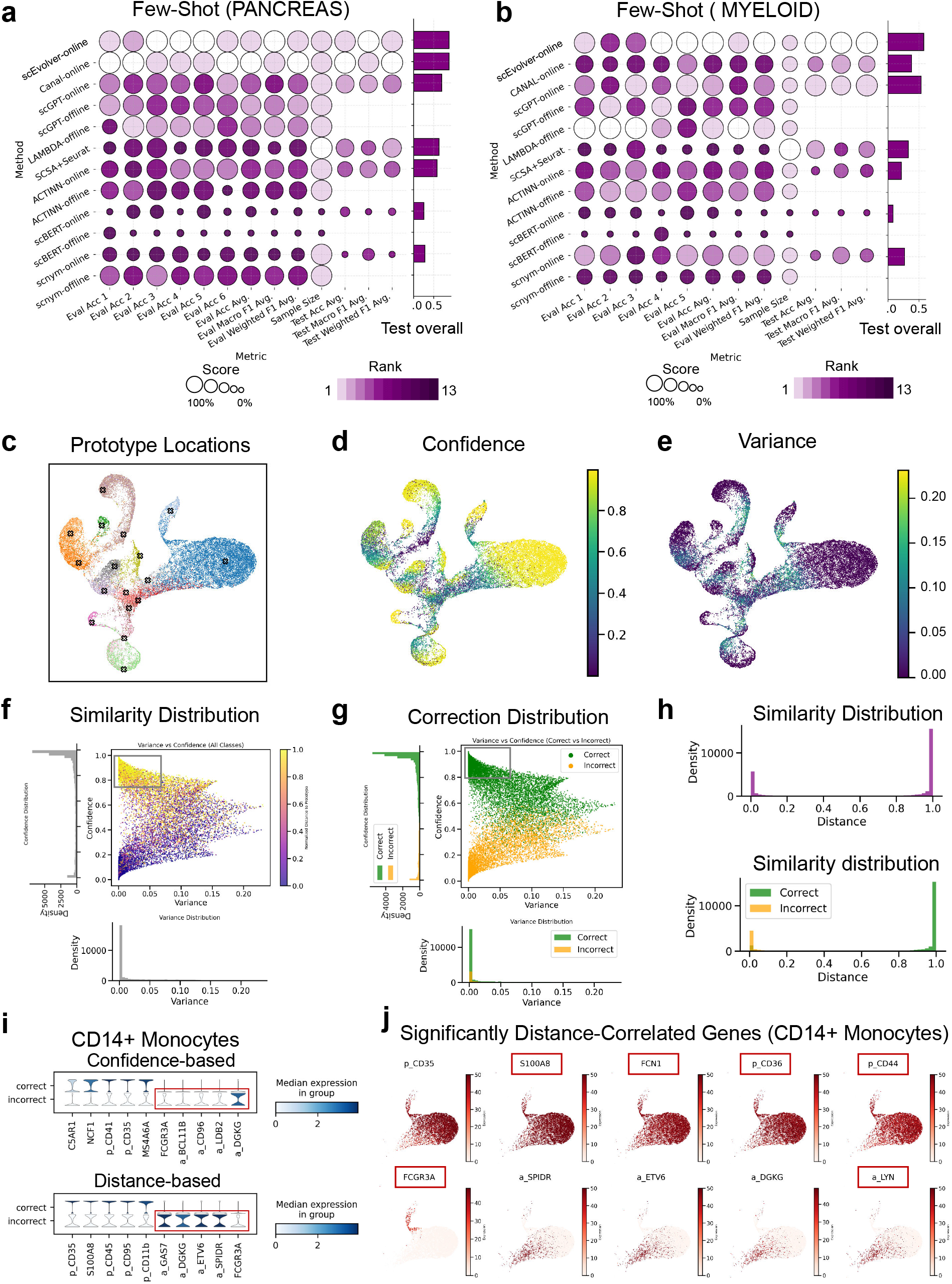
Robust and generalizable single-cell annotation with interpretable cell states. **a, b**, Performance under a few-shot setting with five labeled cells per class. scEvolver mitigates catastrophic forgetting while maintaining robust annotation performance across cell types, ranking first among all benchmarked methods, including offline baselines. Eval ACC n (n = 1–6) shows the accuracy of batch n on the validation set in the retrospective validation experiment. Eval ACC Avg., Eval Macro F1 Avg., and Eval Weighted F1 Avg. represent the overall accuracy, macro-F1 score, and weighted-F1 score across all validation batches, respectively. Sample size indicates the number of samples used. Marker-based methods use the full sample set, while scBERT uses fewer samples due to data filtering during original preprocessing. Other methods have consistent sample sizes. Test ACC Avg., Eval Macro F1 Avg., and Eval Weighted F1 Avg denote the overall accuracy, macro-F1 score, and weighted-F1 score on all held-out test sets, respectively. **c-e**, Distribution of class prototypes (dark ‘×’ markers) and samples in the prediction confidence–variance space. Most prototypes localize to regions of high confidence and low-variance, indicating stable representations. **f, g**, Scatter plots in the prediction confidence–variance space, with point colors indicating sample–prototype similarity (left) and classification correctness (right). Higher confidence and lower variance are associated with greater prototype similarity and improved classification accuracy. **h**, Distributions of similarity scores (top) and classification outcomes (correct versus incorrect; bottom). **i**, Violin plots showing expression distributions of key genes across correctly and incorrectly classified samples. Genes were identified through differential expression analysis (top five upregulated in each group) and correlation analysis with prototype distance (top five positively and negatively correlated genes). **j**, Expression patterns of significantly correlated genes across cell clusters, revealing three monocyte subpopulations—FCGR3A^+^, LYN^+^, and CD14^+^ monocytes—distinguished by marker gene expression.

**Fig. 6.**
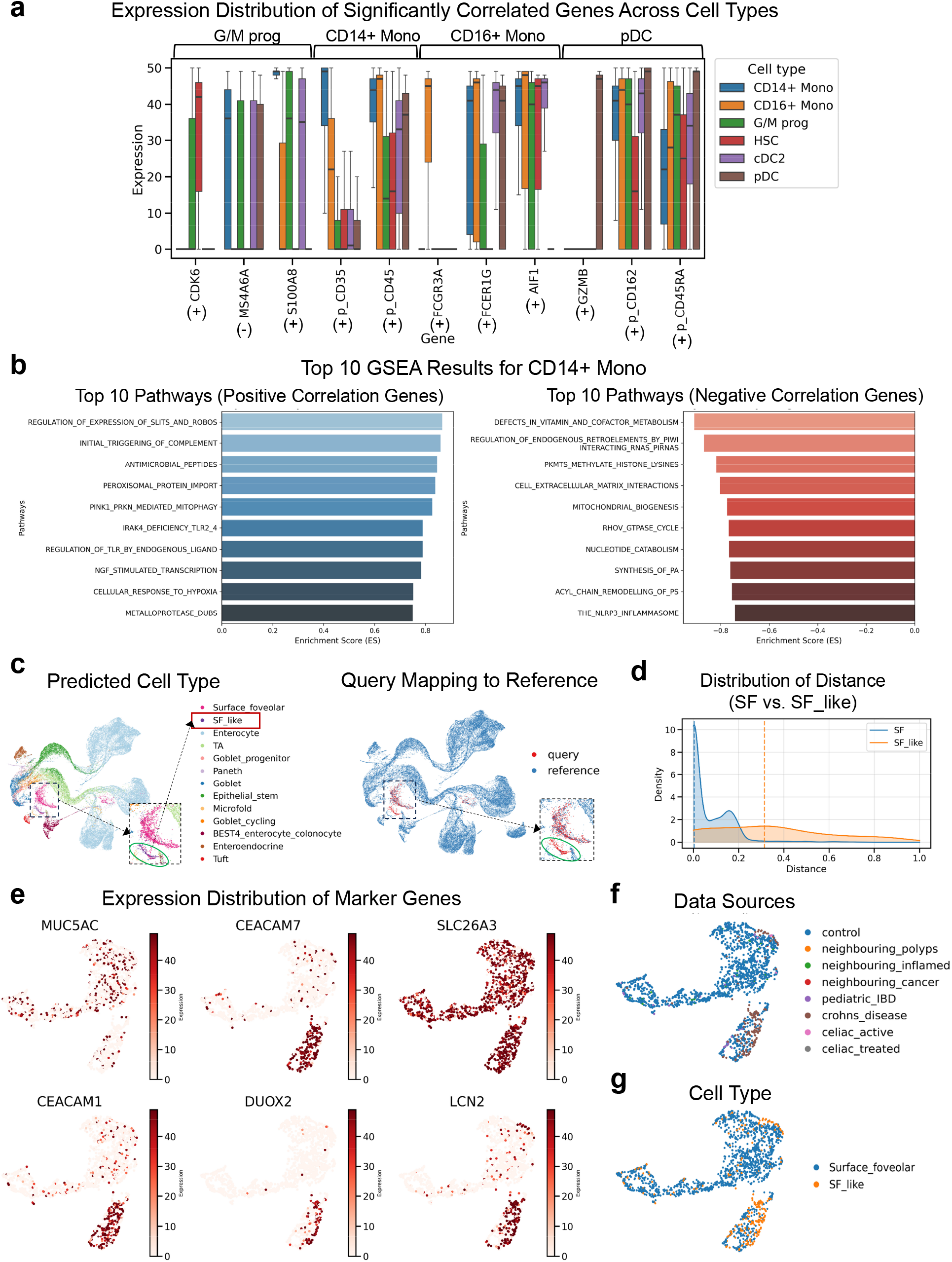
scEvolver reveals prototype distance–associated pathway enrichment and epithelial cell-state transitions in disease contexts. **a**, Expression distribution of genes with high prototype distance correlation across myeloid cell types, including G/M progenitors, CD14+ monocytes, CD16+ monocytes. Most of these genes exhibit strong cell-type specificity simultaneously (54 of 61; supplementary Table S7). **b**, Reactome pathway enrichment of genes significantly correlated with prototype distance in CD14+ monocytes. Positively correlated genes are enriched for canonical innate immune pathways, including complement activation, antimicrobial effector responses, and Toll-like receptor signalling. Negatively correlated genes are enriched for pathways related to inflammasome activation, cytoskeletal regulation, and cell–extracellular matrix interactions. The top ten enriched pathways are shown. **c**, UMAP visualization of disease query cells showing clear separation between SF-like cells and canonical SF cells. **d**, Distribution of normalized distances of SF and SF-like cells to the SF prototype. SF cells exhibit a bimodal distribution with peaks near 0 and 0.18, while SF-like cells show a concentrated distribution centered around 0.3, indicating a measurable deviation from the canonical SF state. **e**, UMAP visualization of established SF lineage marker gene expression. SF-like cells show elevated expression of CEACAM7, CEACAM1, DUOX2, and LCN2, with expression varying smoothly along the embedding manifold. **f-g**, UMAP visualization of cell embeddings showing tissue origin and cell-type identity.

**Fig. 7.**
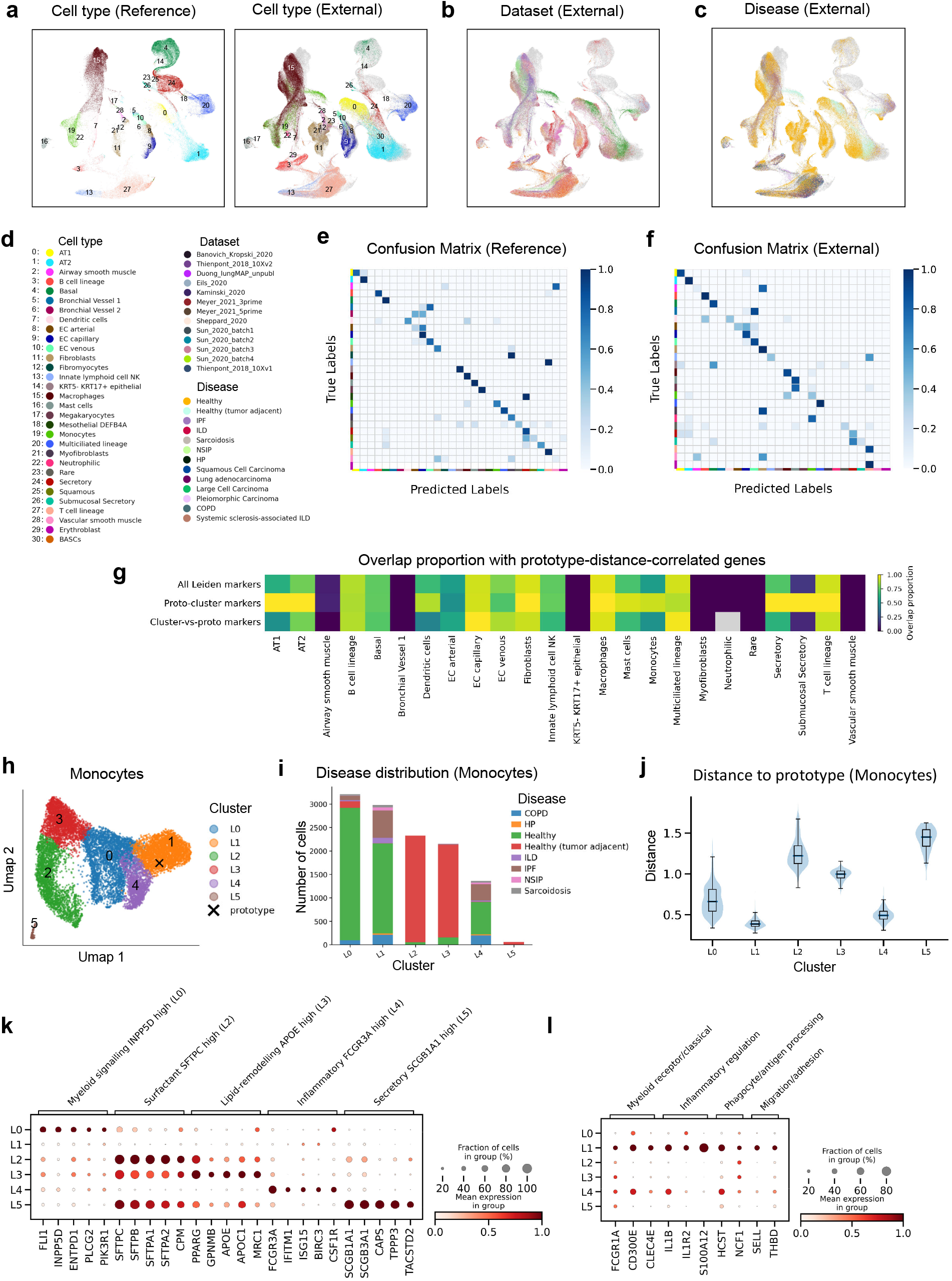
Cross-disease mapping in the HLCA healthy prototype space reveals recurrent aberrant and discordant lung cell states. **a-c**, UMAP visualizations of cells mapped into the prototype-anchored latent space, coloured by annotated cell type in the reference and external datasets (**a**), external dataset source (**b**) and lung disease condition (**c**). **d**, Colour legends for cell types, datasets and disease conditions shown in **a-c. e**,**f**, Normalized confusion matrices showing cell-type prediction performance in the reference validation split (**e**) and external mapped datasets (**f**). Rows indicate true labels and columns indicate predicted labels. The corresponding label legends are shown in **d** “celltype”. **g**, Overlap proportions between prototype-distance-correlated genes and three marker-gene sets across cell types: all Leiden subcluster markers, prototype-containing subcluster markers and markers distinguishing non-prototype subclusters from the prototype-containing subcluster. **h**, UMAP of monocyte-labelled cells coloured by Leiden subcluster. The cross indicates the reference monocyte prototype. **i**, Lung-disease composition of monocyte Leiden subclusters. **j**, Distribution of distances from cells in each monocyte subcluster to the reference monocyte prototype. The centre line indicates the median, box limits indicate the first and third quartiles, and whiskers extend to 1.5 times the interquartile range; violin width represents the density of cells. **k**,**l**, Dot plots showing representative marker genes of monocyte subclusters. Dot size indicates the fraction of cells expressing each gene, and colour indicates mean expression within each subcluster.

## 2 Results

### 2.1 Overview of scEvolver

scEvolver is formulated as a prototype-guided neural network that maps cell expression profiles into a shared embedding space. In this space, each class prototype represents the central tendency of its reference samples and serves as a representative anchor for that cell type Snell et al. [2017]. Unlike static prototypes, these representations are updated online as new data become available, enabling continual refinement of cell-type annotations across sequential single-cell datasets (Fig. 1). Built on a pre-trained foundation model, scEvolver applies parameter-efficient fine-tuning (PEFT) Han et al. [2024] to support continual learning. Training is guided by a prototype-based objective that promotes compact clustering of cells around their corresponding class prototypes in the embedding space, providing an explicit inductive bias for preserving cell-type semantics as new data are incorporated.

To address the highly imbalanced class distributions commonly observed across datasets, scEvolver further incorporates memory prototypes and data replay strategies to accumulate knowledge while mitigating catastrophic forgetting. Memory prototypes preserve class representations learned from earlier reference datasets, while a sample-level memory bank retains hard examples that are periodically replayed during training. Together, these mechanisms stabilize learning and reinforce alignment between incoming samples and their corresponding prototypes, which is particularly beneficial for rare cell types with limited supervision.

Beyond automated annotation, class prototypes also provide an interpretable reference for characterizing within-class cellular variation. In the shared embedding space, cells that deviate from their class prototype reflect local shifts in embedding geometry, capturing continuous variation in cell state. To quantify these shifts, we define a cell–prototype similarity score that relates gene expression to distances from the class prototype (see Methods). This score spans a continuum from prototypical to peripheral cells within a class, highlights prototype-associated genes that contribute to class separation, and supports downstream pathway enrichment analysis of underlying biological processes. By tracking such deviations, scEvolver enables robust outlier detection and the identification of anomalous cell states or disease-associated transitions.

### 2.2 scEvolver continually harmonizes representations across platforms and tissues

Single-cell transcriptomics provides the primary data source for pretraining foundation models, yet continual integration of heterogeneous datasets poses challenges due to platform- and tissue-specific variation, severe class imbalance, and inconsistent cell-type composition. To evaluate scEvolver under this realistic setting, we designed two complementary benchmarks that test performance across platforms and across tissues.

We first evaluated scEvolver on the human pancreas (PANCREAS) Cui et al. [2024] dataset, which aggregates five independent studies spanning nine batches generated by distinct sequencing technologies (Fig. 2a). The first six batches were introduced sequentially to assess continual knowledge accumulation, while the remaining three batches were held out as an independent test set to evaluate generalization. This dataset exhibits pronounced class imbalance and batch-specific cell-type composition (Fig. 2a). For example, ductal cells predominate in Celseq Hashimshony et al. [2012], whereas alpha and beta cells are dominant in Celseq2 Hashimshony et al. [2016] and Fluidigm C1 Xin et al. [2016], respectively. In parallel, we curated a pan-cancer dataset of tumor-infiltrating myeloid cells (MYELOID) Cheng et al. [2021] comprising 54,241 cells across eight cancer types. scEvolver was trained on data from kidney cancer (KIDNEY, 22,984 cells), myeloma (MYE, 5,860 cells), pancreatic adenocarcinoma (PAAD, 2,853 cells), thyroid carcinoma (THCA, 4,423 cells), and uterine corpus endometrial carcinoma (UCEC, 7,505 cells), and evaluated on independent test sets from esophageal carcinoma (ESCA, 6,113 cells), lymphoma (LYM, 615 cells), and ovarian or fallopian tube carcinoma (OV-FTC, 3,888 cells) (Fig. 3a). This split enables systematic evaluation of cross-tissue generalization under substantial tissue-specific heterogeneity. To further characterize heterogeneity across platforms and tissues (referred to as batches), we visualized gene expression profiles using uniform manifold approximation and projection (UMAP)McInnes et al. [2018], with cells labeled by cell type and batch (Fig. 2b–c and Fig. 3b–c). Cells assigned to the same annotated type but originating from different datasets formed distinct, dataset-specific structures rather than overlapping in the embedding space. This pattern reflects substantial technical and biological variation, as well as inconsistent semantic alignment of cell types in the original data. These observations underscore the difficulty of directly integrating heterogeneous scRNA-seq datasets across platforms and tissues without explicit alignment mechanisms. Rather than fixing cell-type composition across batches or tissues, we adopted a class-incremental learning setting in which the number of cell types can increase or decrease during continual learning, while preserving semantic consistency in a shared prototype space. In this online learning setup, scRNA-seq datasets from different sequencing platforms were added incrementally, with training performed only on the newly introduced batches, without revisiting previous data. As new batches were added, cells of the same type from different platforms or tissues aligned in the shared latent space, minimizing platform-specific or tissue-specific batch effects (Fig. 2d-e, supplementary Fig. S3a-b). Class prototypes evolved with incoming data while retaining information from earlier stages, remaining well aligned with cluster embeddings and indicating stable class-level representations. We benchmarked scEvolver against other annotation methods using a retrospective validation strategy Kirkpatrick et al. [2017], in which performance on previously observed datasets was evaluated under the current model parameters after online learning. scEvolver achieved validation performance comparable to offline models trained on individual datasets (supplementary Fig. S4a–b), indicating minimal forgetting of earlier batches. On previously observed pancreas and myeloid datasets, scEvolver achieved average macro F1 scores of 0.9584 and 0.8023, respectively, ranking first among all online approaches (supplementary table S1). Together, these results demonstrate that scEvolver effectively harmonizes representations across sequencing platforms and tissues while mitigating catastrophic forgetting.

### 2.3 scEvolver enables generalizable query mapping and outlier detection

We further evaluated scEvolver on downstream cell annotation tasks, including independent query data mapping, rare cell-type annotation, and outlier detection.

On the PANCREAS dataset, query cells from the held-out test set were mapped into the reference embedding and prototype spaces. Cell embeddings consistently aligned with their corresponding class prototypes, and predicted labels closely matched ground-truth annotations, as visualized by UMAP (Fig. 2f and supplementary Fig. S5a). Quantitative evaluation using bubble and bar charts showed that scEvolver achieved classification performance comparable to CANALWan et al. [2024] on the query test datasets, with the test overall score calculated as the arithmetic mean of accuracy, macro-F1 and weighted-F1 (supplementary Fig. S4a, supplementary Table S1). In addition, latent space clustering achieved a single-cell integration benchmarking (scIB) overall score of 0.91 (overall score = 0.6 × biology conservation + 0.4 × batch correction), ranking first among all evaluated methods (supplementary Fig. S4c, supplementary Table S2). Consistent with these metrics, scEvolver produced more compact cell-type clusters and improved batch mixing in the UMAP space (supplementary Fig. S1 and S2), indicating effective alignment of biological signals while mitigating technical variation. We next assessed cross-tissue generalization on the MYELOID dataset. scEvolver achieved a batch correction score of 0.8546 and a biology conservation score of 0.6095, resulting in a combined score of 0.7075, which exceeded that of the baseline (0.6327; Fig. 3d–e, supplementary Fig. S4d, and supplementary Table S2). In terms of annotation accuracy, scEvolver achieved an accuracy of 0.6544 and a macro F1 score of 0.6332 on the query datasets, ranking second among all evaluated online methods (supplementary Fig. S4b and supplementary Table S3). UMAP visualizations further confirmed that query cells were accurately mapped to their corresponding reference cell types, forming coherent clusters by cell identity (Fig. 3f).

To further examine alignment at the prototype level, we analyzed the distribution of distances between query cells and class prototypes (see Methods). Across both PANCREAS and MYELOID datasets, most query cells showed high similarity to their assigned prototypes and low similarity to other classes, indicating effective alignment within the shared prototype space (supplementary Fig. S6a and S7a). Notably, a subset of cells exhibited overlapping distance profiles across biologically related cell types, such as Macro_C1QC and Macro_LYVE1, or CD3_LAMP3 and cDC2_CD1C. These overlaps reflect shared lineage or functional programs, indicating preserved biological continuity in the latent space, rather than erroneous alignment with unrelated cell types.

To evaluate outlier detection when certain cell types are absent from the reference but appear in query data, we performed controlled exclusion experiments on both the PANCREAS and MYELOID datasets. For the PANCREAS dataset, we manually removed mast, macrophage, and quiescent_stellate cell types from the training set and examined the maximum similarity between query cell embeddings and all class prototypes in the test set (see Methods). Boxplot analysis showed that excluded cell types consistently exhibited low maximum similarity scores (below 0.6), whereas most known cell types had similarity values close to 1 (supplementary Fig. S8a-b, Fig. S6b), enabling clear separation of unseen cell populations. We performed a similar analysis on the MYELOID dataset by excluding pDC_LILRA4, cDC3_LAMP3, and Macro_SPP1 from the training set. UMAP visualization showed that these unseen cell types formed distinct clusters separated from the reference data (Fig. 3g). Consistently, analysis of maximum similarity scores revealed that approximately 75% of unseen cells had values below 0.95, while more than 75% of known cells exceeded this threshold (Fig. 3h). Together, these results indicate that unseen cell types remain distant from all existing class prototypes in the latent space, preventing spurious high-similarity assignments and enabling reliable identification of out-of-distribution cell populations.

### 2.4 scEvolver enables continual mosaic integration for cross-modal single-cell annotation

Most existing single-cell foundation models are primarily trained on scRNA-seq data and therefore have limited capacity to integrate information across multiple modalities. For example, scBERT and the scBERT-based CANALWan et al. [2024] model operate on the gene-token vocabulary defined during pre-training and were not extended to accommodate additional modalities. To overcome this limitation, we adopt a post-training extension strategy that augments a pre-trained foundation model with additional gene tokens representing ATAC and ADT features.

We curated a collection of multimodal datasets derived from bone marrow mononuclear cells (BMMC) Luecken et al. [2021], comprising paired ATAC+RNA and ADT+RNA measurements. In total, the dataset includes 12 multimodal profiles from six distinct sample sources, enabling systematic evaluation of cross-modal representation learning (Fig. 4a–c). During post-training, we employ adversarial learning to encourage modality-invariant representations, thereby reducing technical variation arising from modality-specific biases. At the same time, the introduction of modality-specific tokens allows the model to learn meaningful representations for previously unseen features. Training is guided by a masked token prediction objective, in which masked inputs are reconstructed by predicting the corresponding tokens in the sequence. In the subsequent online learning phase, the model is continuously exposed to incoming data from either ATAC+RNA or ADT+RNA modalities. Under this setting, scEvolver effectively bridges modality differences within individual samples (Fig. 4d) and achieves robust alignment across multimodal datasets (supplementary Fig. S3c–d). UMAP visualizations indicate that the learned latent space supports consistent cell-type alignment across multiple test datasets, while substantially reducing batch-related variation compared with the baseline and other methods (Fig. 4e and supplementary Fig. S9a-b, S1 and S2, supplementary Table S2 and Table S4)

### 2.5 scEvolver improves resistance to catastrophic forgetting and knowledge accumulation

As single-cell datasets continue to expand across sequencing platforms, tissues, and modalities, continual learning provides a promising framework for incrementally updating models without retraining from scratch. A central challenge in this setting is balancing the acquisition of new knowledge with the prevention of catastrophic forgetting. To assess this balance, we systematically evaluated knowledge accumulation across multiple scenarios (Sections 2.2–2.4). To further characterize performance retention, we visualized forgetting curves that track model accuracy on previously observed datasets as new data are introduced (Fig. 4f and supplementary Fig. S10). On the MYELOID dataset, both scNym Kimmel and Kelley [2020] and scGPT Cui et al. [2024] show pronounced performance fluctuations on earlier datasets, characterized by rapid initial forgetting followed by partial recovery. Similar instability is observed for scNym on the BMMC dataset and for ACTINN Ma and Pellegrini [2020] on the PANCREAS dataset. In contrast, ACTINN and scBERT Yang et al. [2022] maintain relatively strong performance on the initial batch but perform poorly on subsequently arriving datasets, indicating overfitting to early data and limited capacity to incorporate new information. Comparable behavior is also observed for scNym and scGPT on the PANCREAS dataset. Overall, scEvolver achieves the most favorable trade-off between knowledge accumulation and resistance to forgetting, indicating a more effective balance between stability and plasticity in continual single-cell annotation.

We further evaluated the contribution of memory replay to mitigating performance fluctuations caused by catastrophic forgetting and to facilitating knowledge accumulation (supplementary Fig. S11a). Models equipped with memory replay (see Methods for replay regime) consistently exhibit reduced performance degradation on previously observed datasets, indicating more effective mitigation of catastrophic forgetting. Analysis of the replay buffer shows that replay retains a higher proportion of samples from earlier datasets (supplementary Fig. S11b). Notably, replayed cells are enriched near cluster boundaries, suggesting that prototype distance– and entropy-based sampling preferentially captures hard-to-classify cells located in ambiguous decision regions (supplementary Fig. S11c–d).

Finally, we compared latent-space organization with and without prototype guidance (supplementary Fig. S11e–f). Incorporating prototype evolution results in smoother state transitions and improved continuity in the latent space. For example, in the region highlighted by the red circle, the model captures a gradual transition from CD14+ to CD16 + monocytes, preserving biologically meaningful relationships across sequentially learned datasets. These results suggest that prototype guidance helps retain cell-state information during continual learning, which we further analyze in Section 2.7.

### 2.6 scEvolver achieves robust, generalizable annotation with fewer labeled samples

In practice, cell-type annotation often relies on a limited number of high-confidence reference cells, typically identified through marker gene expression, expert curation, or targeted experimental validation, rather than exhaustive labeling of all cells. As a result, high-quality labeled examples are typically scarce, particularly for rare or newly characterized cell populations. To assess robustness under label-scarce conditions, we evaluated scEvolver in a few-shot setting, retaining only five labeled cells per class (see Methods). Benchmarking results show that scEvolver effectively mitigates catastrophic forgetting while maintaining robust generalization (Fig. 5a–b). Across independent query test sets, scEvolver achieved strong annotation performance relative to the compared methods, including offline-trained models. On the PANCREAS and MYELOID benchmarks, scEvolver improved macro-F1 scores by 6.1% and 11.6%, respectively, with corresponding gains of 0.3 and 4.5 percentage points in the overall test score (Supplementary Fig. S12a,b; supplementary Table S5 and Table S6). In the latent-space clustering evaluation, scEvolver achieved biological conservation scores of 0.7635 and 0.5696, batch-correction scores of 0.8731 and 0.8849, and overall scIB scores of 0.8073 and 0.6957 on PANCREAS and MYELOID, respectively. These overall scIB scores exceeded those of the best competing methods by 3.5 and 1.0 percentage points, indicating that scEvolver balances biological structure preservation with batch-effect mitigation (supplementary Table S2).

We further examined how annotation performance varies with the amount of available supervision, ranging from 10% to 90% of labeled reference samples. Under this setting, scEvolver was benchmarked against full-batch fine-tuning of the foundation model (see Methods), both with and without PEFT. Radar plot analysis shows that scEvolver maintains stable performance as supervision is progressively reduced (Fig. 4g and supplementary Fig. S13). Across all levels of supervision, scEvolver consistently outperforms full-batch fine-tuning methods, particularly in terms of macro-F1 score, indicating robust performance on imbalanced cell-type distributions. Together, these results demonstrate that scEvolver generalizes effectively under distribution shift while requiring substantially fewer labeled samples in practice.

### 2.7 Prototype distance–correlated genes and pathway enrichment reveal cell-state transitions

In Section 2.5, we showed that the evolutionary prototype strategy in scEvolver captures cell-state transitions while improving representational continuity. Under a Bregman divergence, computing class prototypes as the mean of support sample embeddings yields the optimal cluster representative Banerjee et al. [2005]. Accordingly, the distance to a class prototype provides a principled measure of deviation from the canonical cell-type state. Motivated by this property, we investigated how cell-state transitions are reflected by distances to class prototypes.

We first examined the relationship between the spatial positions of class prototypes in the latent embedding space and the model’s predictive confidence and variance (Fig. 5c–e; see Methods for details). Qualitative analysis shows that most class prototypes (dark ‘×’ markers) are located in regions of high confidence and low variance, indicating stable representations with low predictive uncertainty. To further quantify this relationship, we derived a class probability distribution based on distances to class prototypes. For clarity, we refer to this quantity as “similarity”, rather than explicitly describing it as a “softmax over negative distances”. Figures 5f and 5g show scatter plots in the confidence–variance space, with point colors indicating sample–prototype similarity and classification correctness, respectively. We observe that higher predictive confidence and lower variance are generally associated with greater similarity to the corresponding class prototype and higher classification accuracy. Consistent with this trend, Fig. 5h presents the distributions of similarity scores for correctly and incorrectly classified samples. Statistical analysis reveals a strong correlation between prototype distance and classification correctness (Pearson correlation = 0.7034, p<0.0001). Samples were stratified into correctly and incorrectly classified groups based on their distribution in the prediction confidence–variance space using a confidence threshold (Fig. 5g, see Methods). Differential expression analysis was then performed to identify genes significantly upregulated in each group. In parallel, genes were ranked by the correlation between their expression levels and prototype similarity, yielding the top five positively and negatively correlated genes. Unlike conventional group-wise differential expression analysis, this correlation-based approach does not require predefined sample groups. Group-wise comparisons may compress continuous expression changes into average between-group differences, thereby obscuring subtle cell-state signals. By linking latent-state deviations directly to transcriptional changes, our approach provides an interpretable connection between latent-space deviations and the gene-expression programmes underlying cell-state transitions. The resulting expression patterns of CD14+ Monocyte are shown as violin plots in Fig. 5i. We found that the top 5 genes significantly positively correlated with prototype similarity were highly expressed in correctly classified cells, while the significantly negatively correlated genes were highly expressed in incorrectly classified cells. These observations suggest that prototype similarity is associated with distinct molecular profiles between correctly and incorrectly classified cells, supporting its use as a model-derived indicator of annotation uncertainty and latent representation deviation. Further statistical analysis showed that the top 10 genes identified based on confidence and prototype similarity include overlapping genes such as p_CD35, FCGR3A, and a_DGKG, which are derived from ADT, RNA, and ATAC features, respectively. Figure 5j further visualizes the expression of significantly correlated genes, revealing three monocyte subpopulations distinguished by marker gene expression: FCGR3A^+^ monocytes, LYN^+^ monocytes, and canonical CD14^+^ monocytess, with the marker genes sourced from PanglaoDB Franzén et al. [2019]. Together, these results support the use of prototype distance as a continuous, cell-wise measure of representational deviation without requiring predefined intra-cell-type state groups. Molecular feature–distance correlation analysis further links such latent deviations to interpretable features of RNA expression, ADT protein abundance, and ATAC chromatin accessibility, thereby reflecting transitions in cellular functional states.

Given that prototype distance reflects shifts in gene expression states underlying cell identity, we further investigated whether genes strongly correlated with prototype distance are also specific to particular cell types. To this end, we selected genes with high prototype distance correlation for major myeloid populations, including granulocyte/monocyte (G/M) progenitors, CD14^+^ monocytes, CD16^+^ monocytes, and plasmacytoid dendritic cells (pDCs). Fig. 6a visualizes the distribution of these genes across various cell types. Pairwise statistical analysis revealed that most genes with strong distance correlation also exhibit high cell-type specificity (54 of 61; supplementary Table S7). These findings suggest that prototype distance is closely associated with the expression of key genes defining both cell identity and state, capturing cell-type–specific transcriptional signatures. Accordingly, the similarity between a cell and its class prototype reflects the degree of concordance between the cell’s transcriptional profile and the canonical state of that cell type. Genes showing strong distance correlation with class prototypes were ranked by signed correlation coefficients and subjected to Reactome gene set enrichment analysis (GSEA) Ragueneau et al. [2026], Subramanian et al. [2005]. This analysis identified pathways enriched at the positive and negative extremes of the ranked gene list, with the top 10 results shown in Fig. 6b. Positively correlated genes were enriched for pathways defining canonical innate immune functions of classical CD14^+^ monocytes, including complement activation (INITIAL TRIGGERING OF COMPLEMENT) and antimicrobial effector responses (ANTIMICROBIAL PEPTIDES). Enrichment of Toll-like receptor regulation (REGULATION OF TLR BY ENDOGENOUS LIGAND) and TLR2/4–IRAK4 signaling further highlights the central role of innate immune receptor pathways in classical monocyte activation. In contrast, negatively correlated genes were enriched for pathways associated with inflammasome signaling (THE NLRP3 INFLAMMASOME), cytoskeletal regulation (RHOV GTPASE CYCLE), and extracellular matrix interactions (CELL–EXTRACELLULAR MATRIX INTERACTIONS), consistent with a cell state characterized by heightened inflammatory effector activity and structural remodeling. However, under normal conditions, these pathways are typically suppressed.

To further assess whether scEvolver captures fine-grained, disease-associated cell-state differences, we curated gut data from both healthy individuals and patients, as described in Oliver et al. Oliver et al. [2024]. The resulting small intestine reference dataset comprised 93,891 cells from 116 donors across four tissue regions and 13 datasets. In parallel, a disease query cohort was constructed containing 102,682 single-cell transcriptomes from 43 donors across four tissues and 10 datasets, including gastric and colorectal cancer, ulcerative colitis, Crohn’s disease, and coeliac disease. Through continual cell annotation and query mapping, scEvolver revealed a clear separation between canonical surface foveolar (SF) cells and SF-like cells within the disease query data (Fig. 6c). This pattern is consistent with previous findings by Oliver et al., further confirming the biological validity of the disease-associated state transitions inferred by our model. We next examined the expression patterns of established marker genes within the SF lineage by visualizing their distribution across the learned embedding space, where each point represents an individual cell embedding. SF-like cells showed elevated expression of CEACAM7, CEACAM1, DUOX2, and LCN2, with expression levels varying smoothly along the UMAP manifold (Fig. 6e). This gradual change in expression across the embedding space suggests that the learned representations capture not only discrete cell-type identities but also fine-grained heterogeneity and progressive variation in cellular state. To further quantify the positional relationship between SF-like cells and the canonical reference state, we analyzed the distribution of normalized distances to the SF prototype within the query dataset. SF-like cells exhibited a relatively concentrated distribution centered at a normalized distance of approximately 0.3, indicating a measurable deviation from the canonical SF state. In contrast, SF cells displayed a bimodal distance distribution with peaks near 0 and 0.18, suggesting heterogeneity within the SF population itself (Fig. 6d). Together, these observations support the ability of prototype-based distance metrics to capture continuous variation in cellular states. They further provide an interpretable framework for connecting latent-space deviations with changes in gene-expression programmes and dysregulated pathways during cellular transitions.

### 2.8 Prototype-guided reference mapping reveals common aberrant cell states across diseases

To further evaluate the ability of reference mapping to identify disease-associated cell-state transitions and annotation-discordant populations across datasets, we applied scEvolver to the large-scale Human Lung Cell Atlas (HLCA)Sikkema et al. [2023]. We first sampled 100,000 cells from the HLCA core healthy cohort, covering 29 originally annotated cell types, to construct healthy reference prototypes. To extend this reference and compare cell states between healthy and disease contexts, we mapped 13 external lung datasets onto the healthy reference. These datasets covered 13 lung conditions, including healthy, healthy tumour-adjacent tissue, hypersensitivity pneumonitis (HP), interstitial lung disease (ILD), idiopathic pulmonary fibrosis (IPF), non-specific interstitial pneumonia (NSIP), sarcoidosis, chronic obstructive pulmonary disease (COPD), systemic sclerosis-associated ILD, large-cell carcinoma, lung adenocarcinoma, pleomorphic carcinoma and squamous cell carcinoma, and together comprised 31 cell types (Fig. 7a–d; supplementary Fig. S15a). Label transfer from the HLCA healthy reference achieved 84.47% accuracy in the mapped external datasets and 90.07% accuracy in the reference validation split (Fig. 7e,f, supplementary Fig. S15b-e). Notably, erythroblasts, which were absent from the healthy reference, formed latent representations distinct from existing reference cell types. This indicates that scEvolver can preserve clearly out-of-reference populations in the prototype-anchored latent space rather than forcing them into predefined categories (Fig. 7a). In addition, we observed clear separation between healthy and disease-derived cells within several annotated cell types (e.g. Monocytes and B cell lineage), suggesting that the mapped latent space captures disease-associated deviations from normal cellular states.

To investigate the transcriptional basis of these deviations, we identified prototype-distance-correlated genes for each cell type and performed Leiden Traag et al. [2019] subclustering using three complementary strategies (supplementary meterials). Fisher’s exact test showed that Leiden subcluster marker genes were significantly enriched for prototype-distance-correlated genes in 17 of 24 cell types. Marker genes distinguishing non-prototype subclusters from the prototype-containing subcluster were also enriched for prototype-distance-correlated genes in 13 of 23 cell types (Fig. 7g and the number of overlap genes in supplementary Fig. S16a, supplementary Table S8-12). These overlapping genes simultaneously capture discrete subpopulation heterogeneity and continuous deviations from canonical cell states, highlighting key transcriptional programmes underlying biologically meaningful cell-state variation.

Continual integration of datasets spanning diverse inflammatory, fibrotic and malignant lung conditions further allowed us to examine disease-associated cellular deviations across diseases. Identifying such shared disease-associated cellular states could improve our understanding of common mechanisms underlying lung diseases and facilitate the development of effective treatments. For example, L1 had the smallest mean distance to the reference monocyte prototype (Fig. 7j). It consisted mainly of healthy lung monocytes but also included cells from all disease conditions, indicating a shared reference-like monocyte state. Consistently, L1 expressed genes associated with myeloid immune recognition, inflammatory readiness and leukocyte trafficking, including FCGR1A, IL1B, S100A12 and SELL (Fig. 7l), with APOBEC3A, FCGR1A, IL1B and IL1R2 overlapping normal lung monocyte markers from CellMarker2.0 Hu et al. [2023]. In contrast, prototype-divergent subclusters showed disease-associated programmes. In agreement with previous reports of FCGR3A^+^ monocytes or monocyte-derived macrophage populations in COPD and sarcoidosis Garman et al. [2020], Pei et al. [2022], Zhang et al. [2024], we identified a prototype-divergent inflammatory state with high FCGR3A expression (cluster 4; IFITM1^*high*^, BIRC3^*high*^ and ISG15^*high*^; Fig. 7h–i,k; supplementary Fig. S16b-c). FCGR3A encodes CD16a/Fc*γ*RIIIa, an IgG receptor involved in immune-complex recognition and antibody-dependent inflammatory effector functions. The presence of this cluster in sarcoidosis, COPD and IPF suggests a recurrent inflammatory monocyte programme shared across these lung diseases. We further observed that L2 and L3 were enriched in healthy tumour-adjacent samples and transcriptionally separated from canonical healthy monocyte clusters. Importantly, these healthy tumour-adjacent samples were derived from non-affected surgical lung tissue of lung adenocarcinoma patients Lukassen et al. [2020]. Of these two subclusters, L2 displayed a surfactant-associated epithelial programme marked by high expression of SFTPC, a gene involved in pulmonary surfactant homeostasis (cluster 2; SFTPA1/2^*high*^, SFTPB^*high*^; Fig. 7h–i,k; supplementary Fig. S16b-c). We also identified a monocytes subset expressing lipid clearance and phagocytosis associated genes, one of which (GPNMB) is being investigated as a therapeutic target for squamous cell carcinoma Khan et al. [2021] and other solid tumors Zemp et al. [2026], Baker et al. [2026] (cluster 3; APOE, to play a crucial role in lipid transport and tissue-remodelling; Fig. 7h–i,k; supplementary Fig. S16b-c). Notably, L5 showed the largest distance from the reference monocyte prototype among all monocyte-labelled subclusters (Fig. 7j; one-sided Mann–Whitney U test versus the remaining subclusters, Benjamini–Hochberg-corrected q < 0.05). This cluster was mainly composed of cells from tumour-adjacent samples and highly expressed genes associated with airway secretory defence and epithelial barrier maintenance (cluster 5; SCGB1A1^*high*^ and SCGB3A1^*high*^; Fig. 7h–i,k; supplementary Fig. S16b-c). L5 also expressed TACSTD2, which encodes TROP2, a clinically targetable cell-surface antigen in NSCLC Sands et al. [2025]. Together, these features identify L5 as an epithelial-associated annotation-discordant population within the monocyte-labelled compartment rather than a canonical monocyte state. Overall, this analysis shows that prototype distance can capture key expression programmes underlying subcluster heterogeneity, identify common aberrant cell states across diseases, and reveal annotation-discordant populations.

## 3 Discussion

The rapid expansion of single-cell omics highlights the need for annotation frameworks that are both accurate and scalable as new data continue to accumulate. By incrementally updating class prototypes in a shared embedding space, scEvolver avoids full-model retraining, reducing computational costs and privacy risks. Our results show that scEvolver effectively mitigates catastrophic forgetting while retaining strong sensitivity to rare cell populations. In addition, the framework unifies reference atlas construction and query mapping within a single system, providing an interpretable link between cell positions in the embedding space and cell-type–specific gene signatures.

While scEvolver performs effectively across multiple modalities, our approach currently assumes that missing modalities are represented by zeros. In some cases, this simplification may not fully capture the biological complexity underlying missing data. Future work could explore imputation strategies to better model missing modalities and improve performance in more complex biological settings. In addition, although scEvolver demonstrates strong performance across the evaluated settings, the current study considers a limited set of practical scenarios. As single-cell technologies continue to advance, more diverse and complex settings are expected to emerge, including novel combinations of multimodal data and increasingly heterogeneous data streams. Incorporating more flexible or dynamic network architectures may further enhance the adaptability and scalability of the framework. With respect to data modalities, scEvolver currently integrates transcriptomic, proteomic, and chromatin accessibility measurements. Extending the framework to additional omics modalities, such as spatial transcriptomics and other spatially resolved technologies, represents an important direction for future research.

Nevertheless, scEvolver not only provides an automated framework for continual cell annotation but also offers a scalable approach to single-cell analysis with direct relevance to disease research and precision medicine. By enabling continual integration of new data, it supports the construction of dynamic cellular references that capture biological variation across conditions and stages. More broadly, this framework provides a foundation for modeling disease state transitions and therapeutic responses, helping advance virtual cell research in cellular heterogeneity.

## 4 Methods

### 4.1 The scenario of single-cell annotation

Single-cell annotation can be formulated under different learning paradigms, depending on data accessibility, label availability, and the ability to accumulate knowledge. Here, we distinguish several representative scenarios. As the incremental datasets are introduced sequentially,

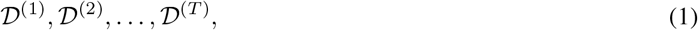

where each dataset

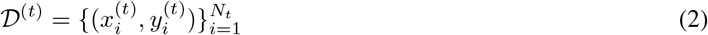

consists of cell expression profiles 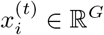 and corresponding cell-type labels 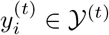. Let *f*_*θ*_ denote a parametric model with parameters *θ*.

#### Full-batch learning

Full-batch learning assumes that all datasets are simultaneously accessible:

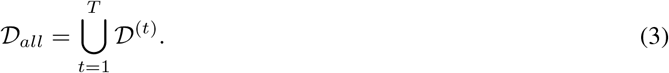

A single model is trained by minimizing the empirical risk over the aggregated dataset:

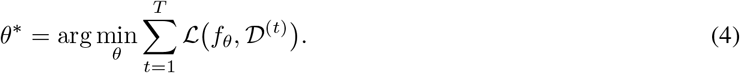

This setting represents an idealized upper bound in which neither temporal ordering nor data access constraints are imposed.

#### Offline learning

In offline learning, only the dataset available at the current stage, denoted as *D*^(*k*)^, is accessible. Models trained at different stages are independent, with no information shared between stages. The learning objective is:

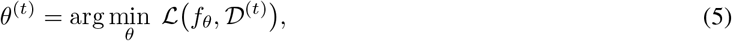

where

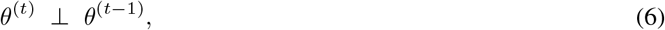

meaning that the model parameters are reinitialized at each stage. This setting is commonly used in scenarios where data sharing is restricted, or historical data cannot be accessed due to privacy, storage, or policy limitations.

#### Continual learning

In continual learning, datasets arrive sequentially and previously observed data are no longer accessible. Model parameters are updated incrementally according to

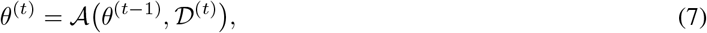

The objective is to preserve performance on previously encountered data while adapting to newly arriving datasets, which can be expressed as

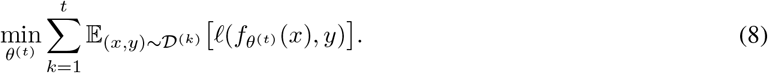

Continual fine-tuning represents a special case of continual learning in which the model is initialized from a pretrained foundation model 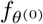.

#### Few-shot offline learning

Few-shot offline learning assumes that only a small labeled subset is available for each dataset:

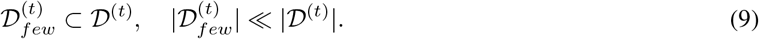

Models are trained independently for each dataset according to

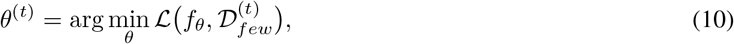

with no parameter sharing across datasets.

#### Few-shot online learning

Few-shot online learning combines sequential data arrival with limited supervision. Model parameters are updated incrementally according to

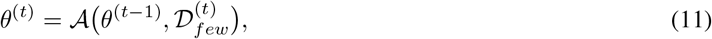

where only a small number of labeled samples are available at each learning stage and previously observed data are not accessible. This setting simultaneously challenges the model in terms of data accessibility, label scarcity, and its ability to accumulate knowledge.

### 4.2 The continual learning framework of scEvolver

scEvolver is a prototype-based continual learning framework for single-cell annotation under non-stationary data streams. The framework is designed to learn stable and discriminative representations by combining parametric classification with memory-augmented, prototype-based supervision.

#### 4.2.1 Representation Learning with PEFT

ScEvolver adopts the scGPT foundation model as the representation learning backbone to encode high-dimensional single-cell gene expression profiles into contextualized latent embeddings. scGPT is pretrained on large-scale single-cell data and captures rich biological semantics across diverse cellular contexts. However, directly fine-tuning all model parameters is computationally expensive and lead to catastrophic forgetting.

To address this limitation, scEvolver employs a PEFT strategy based on low-rank adaptation (LoRA) Hu et al. [2022], enabling efficient and flexible adaptation of the foundation model. During PEFT, the original parameters of the pretrained model are kept frozen, thereby preserving knowledge acquired during pretraining.

The model architecture consists of three main components: a tokenizer, an encoder, and a projector. The encoder is built from 12 stacked Transformer layers Vaswani et al. [2017], each including a multi-head self-attention module, layer normalization, and a feed-forward network (FFN).

In a standard Transformer architecture, the feed-forward network block can be written as

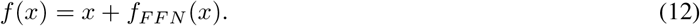

where the FFN typically consists of a linear transformation followed by a non-linear activation function. The linear transformation can be expressed as

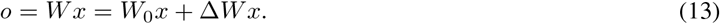

where 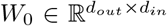 denotes the frozen weight matrix of the pretrained backbone, and Δ*W* represents the task-specific parameter update during fine-tuning.

To enable parameter-efficient adaptation, scEvolver replaces the linear layer in the FFN block with a mixture-of-experts (MoE) module Dou et al. [2024]. During training, the backbone parameters *W*_0_ remain frozen to preserve pretrained biological knowledge, and only the additional parameters are updated. For a MoE layer with *N* experts 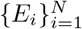, the forward computation is given by

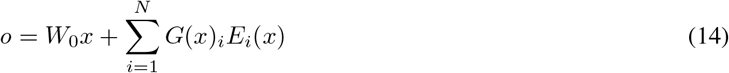

where *G*(·) denotes a routing function defined as

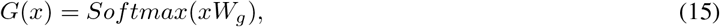

and *W*_*g*_ is the trainable parameter matrix of the routing network.

Instead of introducing a full set of parameters comparable to those in the FFN, LoRA has been shown to provide an effective and parameter-efficient adaptation strategy during supervised fine-tuning (SFT). Specifically, the expert-specific weight update 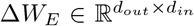 is decomposed as Δ*W*_*E*_ = *QP*, where 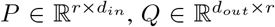, and *r* ≪ min(*d*_*in*_, *d*_*out*_). This low-rank factorization significantly reduces the number of trainable parameters while maintaining task adaptability. By combining the routing mechanism with low-rank experts, the forward computation of the MoE layer is given by

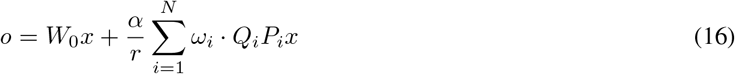

Where *ω*_*i*_ = *G*(*x*)_*i*_ denotes the routing weight of the *i*-th expert and *α* is a scaling hyper-parameter that controls the contribution of the adaptation module.

#### 4.2.2 Prototype-based representation

Prototypical representation address few-shot regimes and outlier detection by introducing a simple inductive bias, in which cells of the same type are assumed to cluster around a shared prototype in a learned embedding space. Formally, given a labeled support set 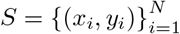, where *x*_*i*_ ∈ ℝ^*D*^ denotes an input feature vector and *y*_*i*_ ∈ *{*1, …, *K}* is the corresponding class label, an embedding function *f*_*ϕ*_ : ℝ^*D*^ → ℝ^*M*^ maps each input into an *M* -dimensional latent space. For each class k, the prototype *p*_*k*_ ∈ ℝ^*M*^ is computed as the mean of the embedded samples belonging to that class:

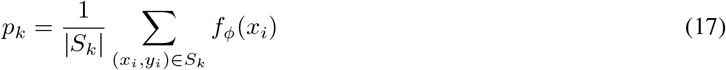

where *S*_*k*_ denotes the subset of samples with label *k*. Under a regular Bregman divergence, the optimal cluster representative (the Bregman centroid) corresponds to the mean of the associated latent representations; a formal derivation is provided in Appendix A. Given a distance function *d*(·,·), such as Euclidean distance, prototypical networks estimate the probability that a query sample *x* belongs to class *k* by comparing its embedding with class prototypes in the latent space:

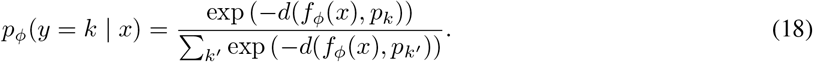

#### 4.2.3 Dual-level Memory Augmentation

##### Memory-augmented prototype

To preserve class-level representations during continual learning, the model maintains a prototype memory buffer *M*_*p*_(*t*), which stores historical class prototypes from earlier stages. At each learning stage *t*, the prototype representation for class *c* consists of the current prototype 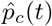 combined with the historical prototypes 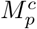. Accordingly, the memory-augmented prototype set is defined as

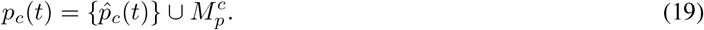

As new data are incrementally added during continual learning, the prototype memory buffer *M*_*p*_(*t*) is updated to include the latest class prototypes, allowing the model to retain previously learned representations while adapting to new data.

##### Memory-augmented sample replay

In parallel, each class maintains a replay sample set *M*_*c*_(*t*). During training, mini-batches are constructed by mixing samples from the current dataset with replayed samples drawn from *M*_*c*_(*t*), ensuring continued exposure to previously observed data distributions. This replay mechanism provides direct supervision signals that help preserve decision boundaries learned at earlier stages. Replayed samples are selected using a hard-sample priority strategy, in which samples are ranked based on prediction entropy and their distance to the current class prototype. Samples with higher entropy and larger prototype distance are preferentially replayed. Empirically, such samples tend to lie near decision boundaries and are more challenging to classify. Compared with raw sample replay alone, prototype memory provides complementary global structural guidance in the latent space. Together, replayed samples from *M*_*c*_(*t*) and historical prototypes stored in *M*_*p*_(*t*) enable scEvolver to balance plasticity and stability during continual learning.

#### 4.2.4 Memory-augmented Prototypical Proxy Loss (MAPPL)

In MAPPL, the model optimizes *f*_*θ*_ to project a sample 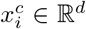 from class *c* close to its memory-augmented class prototype *p*_*c*_(*t*) in the latent space, while simultaneously encouraging separation from prototypes of other classes.

##### Similarity and Dissimilarity Sets

For each class *c*, we define the set of samples belonging to class *c* in the current mini-batch as *D*_*c*_ = *{*(*x*_*i*_, *y*_*i*_ = *c*) ∈ *D}*. Given an anchor sample 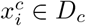, we construct a *similarity set* 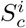, which consists of two components: (i) embeddings of other samples from the same class within the batch, and (ii) a memory-augmented prototype set *p*_*c*_(*t*), which includes the current class prototype and historical prototypes stored in the prototype memory buffer 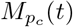. Formally, the similarity set is defined as

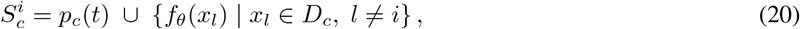

where the memory-augmented prototype set is given by

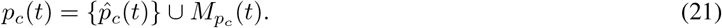

These reference representations encourage embeddings of the same class to align consistently in the latent space. To enforce inter-class separation, we further define a *dissimilarity set* 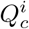, which consists of the memory-augmented prototype set *p*_*c*_(*t*) together with the anchor embedding itself:

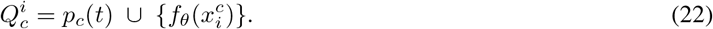

Embeddings from other classes are encouraged to remain distant from the reference representations in 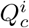, providing repulsive signals that separate class *c* from the rest of the embedding space.

##### MAPPL Objective

A schematic illustration of the distance-based class probability computation and the MAPPL objective is shown in supplementary Fig. S14. MAPPL aims to correctly classify instances of class *c*, while preventing instances from other classes from being assigned to class *c*. This objective is formalized by the following loss function:

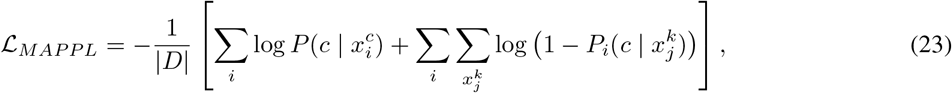

where 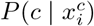 denotes the probability that an instance 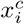 from class *c* is correctly classified as class *c*, and 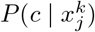 denotes the probability that an instance 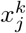 from a different class *k*≠ *c* is incorrectly assigned to class *c*.

Next, we derive the expression for the classification probability. We assume that the distance between an embedded instance and a class prototype is measured by a Bregman divergence, with the squared Euclidean distance as a canonical choice. Specifically, according to Eq. (18), given the Bregman divergence 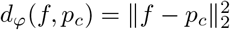, the probability of assigning an instance with embedding *f* (*x*) to class *c* is defined as:

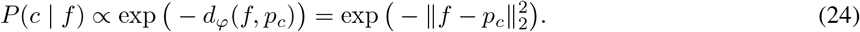

To align with the use of a similarity-based formulation rather than a distance-based one, we exploit the fact that, when embeddings are *ℓ*_2_-normalized, the squared Euclidean distance is equivalent (up to a constant) to cosine distance. Specifically, for unit-normalized vectors *f* and *p*_*c*_, we have:

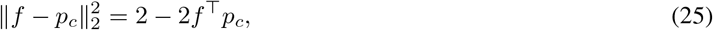

Substituting this relation into the exponential distance-based formulation yields a similarity-based expression:

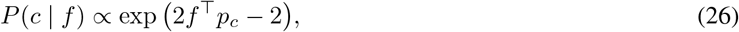

where the constant term does not affect the normalized posterior distribution. Accordingly, the class posterior probability for an instance 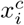 with embedding 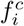 is computed using a softmax over cosine similarities to all class prototypes. Given *K* classes, we define:

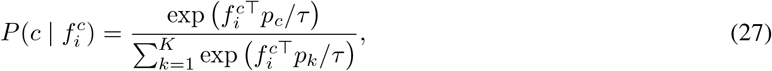

where *τ* is a temperature parameter controlling the sharpness of the distribution.

For the similarity set, we define 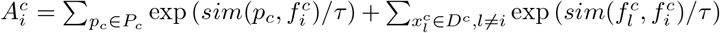, the posterior probability in Eq. (27) can be expressed as:

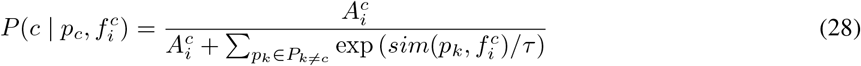

where *P*_*c*_ denotes the set of prototypes associated with class *c*. Similarly, for the dissimilarity set, we define 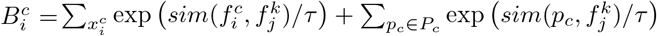, the posterior probability is defined as:

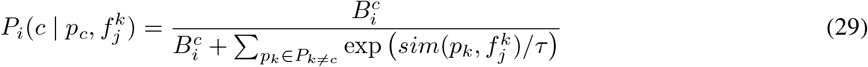

where *P*_*k*≠*c*_ contains the prototypes of all classes other than *p*_*c*_. Finally, we define the expected posterior probabilities for the similarity and dissimilarity sets associated with an anchor instance 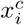 as:

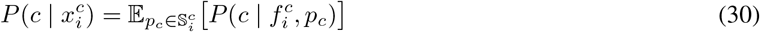

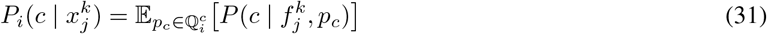

#### 4.2.5 Expandable classification head

In class-incremental learning, newly observed data *X*_*t*_ may introduce additional cell types or categories that were absent in previous training stages. To accommodate the expansion of the label space, we design the classification head to be parameter-expandable, allowing new classes to be incorporated without modifying the feature extractor.

Let 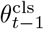 denotes the parameters of the classification head corresponding to previously learned classes, and let 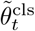 represents the parameters associated with newly introduced classes at step *t*. The new parameters are randomly initialized and concatenated with the existing ones to form the expanded initialization

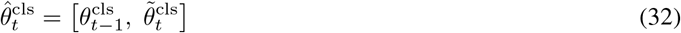

Starting from 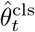, the classification head parameters 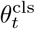 are optimized using the current training data *X*_*t*_ together with the retained reference set *M*_*c*_(*t*).

### 4.3 Model training of scEvolver

#### Cross-platform and cross-tissue training

ScEvolver enables continual cell-type annotation across heterogeneous single-cell datasets spanning multiple sequencing platforms and tissues. The model comprises a tokenizer, a PEFT-based encoder, a decoder, and an expandable classification head. These components are iteratively optimized by minimizing the following joint objective:

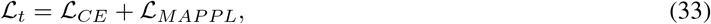

where the loss is evaluated over the current training data *X*_*t*_ together with replay samples *M*_*c*_(*t*). Here, ℒ_CE_ denotes the standard cross-entropy loss, and ℒ_MAPPL_ denotes the memory-augmented prototypical proxy loss.

#### Cross-modality training

For cross-modality prediction tasks, the foundation model must acquire representations for newly introduced feature tokens. To accommodate this requirement, we concatenate the new tokens with previously learned tokens and adopt a post-pretraining adaptation strategy. Specifically, pretrained embeddings are retained for existing tokens, whereas embeddings corresponding to newly introduced tokens are randomly initialized. The model is subsequently optimized using the same continual learning procedure as in the cross-platform and cross-tissue setting.

#### Prototype update

Class prototypes are updated at the end of each training epoch. Given the samples observed at the current continual learning stage *t*, including the current dataset *D*_*t*_ and the replay memory *M*_*c*_(*t*), we first compute their latent representations as 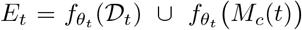. As the data observed at each stage can be considered stochastic samples from the underlying class distribution, directly updating prototypes using the current batch may lead to unstable estimates. To obtain robust and temporally smooth prototype representations, we adopt a high-momentum update rule. Specifically, for each class *c*, the prototype is updated as:

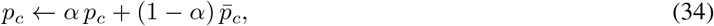

where

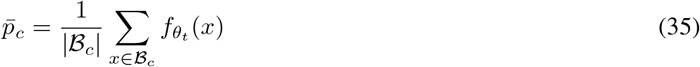

denotes the class-wise mean embedding computed from the current representations, ℬ_*c*_ is the set of samples belonging to class *c*, and *α* ∈ [0, 1] is a momentum coefficient controlling the update rate.

### 4.4 Benchmarking settings

#### Benchmark methods

We selected baseline methods covering marker-based approaches, traditional machine learning, deep learning, and large pre-trained foundation models for single-cell annotation. As these methods are originally designed for offline training, we adapted their implementations to enable evaluation in an online setting. Because they are not developed for continual learning, the number of output classes in their classification heads must be fixed. To allow evaluation across all cell types, we therefore set the output dimension to the maximum number of classes observed across datasets for all baselines. A detailed description of the benchmark methods can be found in the supplementary materials.

#### Classification metrics

Single-cell datasets are often characterized by severe class imbalance and the presence of rare cell types. Rather than discarding minority classes using heuristic thresholds, we retain all annotated cell types in the original datasets and adopt evaluation metrics that explicitly account for class imbalance. Specifically, we use *accuracy* to assess overall classification accuracy and *macro-F1* to calculate the F1 score for each class independently, averaging them while treating all classes equally regardless of their frequency. *Weighted F1*, on the other hand, averages the F1 score for each class, weighted by the number of true instances for each class. The detailed definitions of these metrics can be found in the supplementary materials.

#### Clustering metrics

We evaluated clustering performance using the scIB benchmarking framework, which jointly assesses batch correction and biological conservation. Batch correction performance was quantified using the batch-average silhouette width (ASW) and graph connectivity, whereas biological conservation was assessed using normalized mutual information (NMI), adjusted Rand index (ARI), and cell-type ASW. For each evaluation category, the corresponding metrics were aggregated by averaging to obtain a batch correction score *S*_batch_ and a biological conservation score *S*_bio_. To summarize overall clustering performance while emphasizing preservation of biological structure, we define an overall clustering score as:

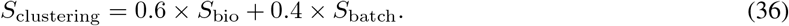

#### Few-shot settings

For each dataset, only five labeled samples per class are retained for training, and classes with fewer than five samples are excluded. This setting is designed to assess the robustness and generalization capability of the model under conditions of severely limited annotated data and constrained computational budgets. To reduce overfitting in this low-sample regime, we employ an early stopping strategy during training. Following preliminary tuning, the batch size is set to 8, and the maximum number of training epochs per dataset is capped at 1000. Early stopping is applied with a patience of 50 epochs, and class prototypes are updated every 10 epochs.

### 4.5 Distance-correlated genes and pathway enrichment analysis

Visualization in the learned embedding space reveals that cells belonging to the same class do not collapse to a single point in UMAP projections. Instead, they exhibit structured spatial dispersion, indicating non-negligible intra-class variation. Such variation likely reflects continuous differences in cellular states within a given cell type. For each cell, we quantify its association with each cell type using a prototype-based similarity score, referred to as *similarity*. This score is defined as the softmax over negative Euclidean distances to all class prototypes (Section 4.2.2, Eq. 18). The pairwise distances and similarities are computed as follows:

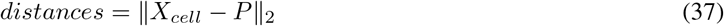

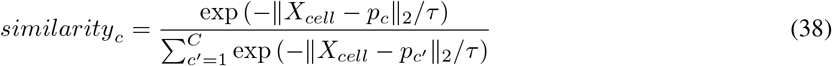

Here, *X*_*cell*_ denotes the embedding of a cell, *p*_*c*_ is the prototype of class *c, C* is the total number of classes, and *τ* is a temperature parameter controlling the sharpness of the similarity distribution (set to *τ* = 0.5 in our experiments).

### 4.6 Confidence and variance estimation

To quantify predictive uncertainty, we adopt a probability-based estimation strategy using Monte Carlo dropout. Specifically, dropout layers remain active during inference, introducing stochasticity through multiple forward passes to obtain class probability predictions. For each sample, we then extract the predicted probabilities of its ground-truth class, denoted as 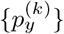. The mean of these probabilities is used as a measure of predictive confidence, whereas their variance reflects predictive uncertainty.

Following Karin *et al*. Karin et al. [2024], samples with high confidence and low variance correspond to predictions that are consistently aligned with the assigned labels. Samples with high variance form an intermediate group with ambiguous predictions. In contrast, samples with low confidence and low variance indicate disagreement between model predictions and labels, and may include mislabeled instances. Based on these properties, we adopted a dataset-dependent approach to quantify sample difficulty. In the Results section, samples were partitioned according to their distribution in the predictive confidence–variability space, with those exhibiting predictive confidence greater than 0.8 classified as correct (easy-to-learn) and those with predictive confidence below 0.2 classified as incorrect (hard-to-learn).

### 4.7 Pathway enrichment analysis

Pathway enrichment analysis was performed using the Reactome database Ragueneau et al. [2026] within a GSEA framework. Reactome pathways were obtained from MSigDB Liberzon et al. [2011], and duplicate gene–pathway mappings were removed. Genes were ranked according to their associated statistics: log fold changes were used for differential expression analyses, and correlation coefficients were used for distance-correlation analyses. Enrichment analysis was conducted using the decoupler implementation of GSEA, yielding pathway-level enrichment scores and corresponding significance estimates.

## 5 Data availability

The original datasets were obtained from the Gene Expression Omnibus (GEO) database. The PANCREAS dataset includes data from Baron (GSE84133), Muraro (GSE85241), Xin (GSE81608), Segerstolpe (E-MTAB-5061), and Lawlor (GSE86473). The processed human pancreas dataset was retrieved from https://github.com/bowang-lab/scGPT. The MYELOID dataset is publicly available under accession number GSE154763 Cheng et al. [2021]. The BMMC dataset was obtained from GEO under accession number GSE194122 and includes two sequencing modalities Luecken et al. [2021]. The small intestinal datasets for the experimental and control groups were obtained from https://gutcellatlas.org/pangi.html Oliver et al. [2024] and were preprocessed using an in-house pipeline, as described in the supplementary materials. The Human Lung Cell Atlas (HLCA) dataset used in this study is publicly accessible through CELLxGENE and contains raw and normalised count matrices, integrated embeddings, cell-type annotations, and clinical and technical metadata: https://cellxgene.cziscience.com/collections/6f6d381a-7701-4781-935c-db10d30de293 Sikkema et al. [2023]. The external datasets used in this study are listed in the supplementary Table S13 and Table S14.

## 6 Code availability

Code is available at https://github.com/AI-HPC-Research-Team/scEvolver.

## 7 Acknowledgments

This work was supported by the Guangdong Science and Technology Programme under Grant 2024B0101010003.

## 8 Supplementary Materials

The supplementary figures referenced in this manuscript are provided in the Supplementary Materials file.

## 10 Appendix A

A regular Bregman divergence *d*_*φ*_ is defined as:

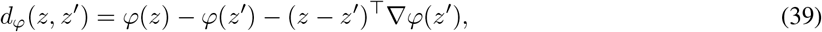

where *φ* is a differentiable, strictly convex function of the Legendre type. Any regular exponential family distribution *p*_*ψ*_(*z* | *θ*) can be expressed in terms of a uniquely determined regular Bregman divergence *d*_*φ*_(·, ·) as

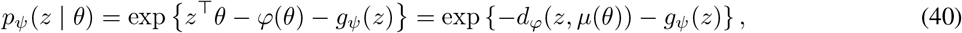

where *µ*(*θ*) = ∇*ψ*(*θ*) represents the mean parameter, and *g*_*ψ*_(*z*) is a base-measure term that does not depend on *θ*. Next, consider an exponential-family mixture model with parameters 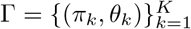:

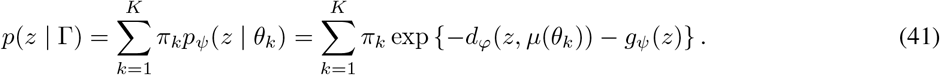

To calculate the posterior probability of assigning an unlabeled sample *z* to class *k*, we apply Bayes’ rule:

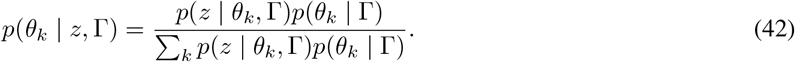

Using the prior *π*_*k*_ = *p*(*θ*_*k*_), we substitute it into the above equation, yielding:

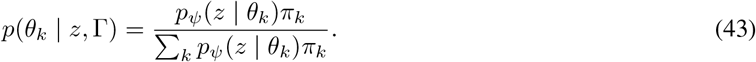

Substituting the expression for *p*_*ψ*_(*z* | *θ*) from Eq. (40) into this, we obtain a form based on the Bregman divergence. Since *g*_*ψ*_(*z*) does not depend on the class index *k*, it cancels out in the normalization, resulting in:

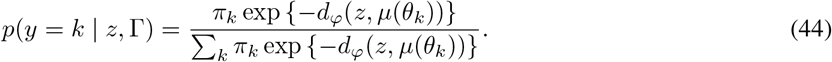

For an equally weighted mixture model (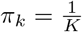 for all *k*), this simplifies to a softmax over the negative distances to the prototypes of each class, which is equivalent to the classification rule used in Prototypical Networks (Eq. (18)), where *z* = *f*_*θ*_(*x*) and *c*_*k*_ = *µ*(*θ*_*k*_).

