## Supplemental Material 1 for "Prototype-based continual cell-type annotation reveals cellular state transitions in expanding single-cell atlases"

A PREPRINT

July 25, 2026

### 1 Supplementary Notes

#### 1.1 Baseline methods

**SCSA** Cao et al. [2020] is a marker-based automated cell type annotation method that assigns cell identities at the cluster level. In our evaluation, cell clusters were first generated using the Seurat Stuart et al. [2019] clustering pipeline, after which SCSA annotated each cluster based on a scoring model that integrates differentially expressed genes with confidence-weighted marker information from curated databases and user-defined marker sets. By leveraging Seurat-derived clusters, SCSA provides a fully automated annotation workflow without manual curation.

**ACTINN** (Automated Cell Type Identification using Neural Networks)Ma and Pellegrini [2020] is a supervised neural network–based method for cell type annotation in single-cell RNA sequencing data. Unlike traditional annotation pipelines that rely on unsupervised clustering followed by manual marker-based labeling, ACTINN directly learns cell type–specific patterns from labeled reference datasets. The model consists of a feedforward neural network with three hidden layers and is trained on datasets with predefined cell type annotations. Once trained, the learned parameters are used to predict cell types in new datasets. ACTINN has been demonstrated to provide fast and accurate cell type predictions across multiple datasets, including mouse and human immune cell atlases, making it a commonly used supervised baseline for scRNA-seq annotation tasks.

**LAmbDA** (Label Ambiguous Domain Adaptation)Johnson et al. [2019] is a transfer learning framework designed for cross-dataset and cross-species cell type annotation in single-cell RNA-seq data. The method jointly addresses batch effect correction, label ambiguity, and cell type mapping by training models on multiple datasets simultaneously, even when subtype labels are inconsistent across sources. LAmbDA is model-agnostic and has been implemented using feedforward neural networks with ensemble strategies, enabling robust subtype-level annotation across heterogeneous datasets. Owing to its ability to integrate multiple references and handle ambiguous labels, LAmbDA serves as a representative transfer learning–based baseline for evaluating cross-dataset cell type annotation performance.

**scNym** Kimmel and Kelley [2020] is a semi-supervised neural network–based method for transferring cell type annotations across single-cell genomics experiments. The model integrates labeled reference data with unlabeled target data to learn shared representations of cell identity, enabling robust annotation transfer across datasets with biological and technical differences. scNym employs an adversarial training strategy to encourage domain-invariant representations and supports the integration of multiple reference and target datasets.

**scBERT** Yang et al. [2022] scBERT is a pretrained transformer-based model for single-cell RNA-seq analysis that adapts the bidirectional encoder representations from transformers (BERT) framework to gene expression data. Through large-scale self-supervised pretraining on unlabelled scRNA-seq datasets, scBERT learns latent gene–gene interaction patterns and generalizable cellular representations. The pretrained model can then be fine-tuned in a supervised manner for cell type annotation on user-specific datasets. By explicitly modeling contextual gene relationships and mitigating batch effects through pretraining, scBERT has demonstrated strong performance in cell type annotation, robustness across batches, and improved interpretability, making it a representative deep learning–based baseline for single-cell annotation.

**scGPT** Cui et al. [2024] is a large-scale foundation model for single-cell biology built on a generative pretrained transformer architecture. It is trained on a diverse repository of over 33 million single-cell profiles, enabling the model to learn generalizable representations of genes and cells across datasets. By drawing analogies between language modeling and cellular gene expression, scGPT captures rich biological semantics through large-scale pretraining. After pretraining, scGPT can be adapted via transfer learning to support a wide range of downstream tasks, including cell type annotation, batch integration, multi-omic integration, perturbation response prediction, and gene network inference. Owing to its scale and pretrained representations, scGPT serves as a strong baseline for evaluating annotation performance under heterogeneous and large-scale single-cell settings.

**CANAL** Wan et al. [2024] is a continual learning method for cell type annotation in single-cell RNA-seq data. It sequentially fine-tunes a pretrained single-cell language model on newly arriving annotated datasets and updates the classifier when new cell types are observed. To alleviate catastrophic forgetting, it uses an example bank and representation knowledge distillation. In our evaluation, CANAL was used as a continual annotation baseline.

#### 1.2 Data preprocessing

Gene expression data were preprocessed using the Scanpy Python library Wolf et al. [2018]. Briefly, low-quality genes and cells with low total counts were optionally filtered. Expression values were then normalized on a per-cell basis by scaling total counts to a fixed value, followed by an optional log1p transformation to stabilize variance. Highly variable genes (HVGs) were identified using the seurat_v3 method Stuart et al. [2019] implemented in Scanpy, with optional batch-aware selection when batch information was available, and only HVGs were retained for downstream analyses. For models requiring discrete inputs, normalized expression values were further transformed using quantile-based binning. Specifically, non-zero expression values within each cell were divided into a fixed number of bins based on empirical quantiles, while zero values were assigned to a separate category. All preprocessing steps were applied consistently across datasets to ensure comparability in downstream evaluations.

##### 1.3 Classification metrics

Let $C$ denote the total number of classes and $n_c$ the number of samples belonging to class $c$, with

$$N = \sum_{c=1}^{C} n_c \tag{1}$$

denoting the total number of samples. For each class $c$, let $\mathrm{TP}_c$, $\mathrm{FP}_c$, and $\mathrm{FN}_c$ denote the numbers of true positives, false positives, and false negatives, respectively.

*Accuracy.* Accuracy is defined as the proportion of correctly classified samples:

$$\mathrm{Accuracy} = \frac{1}{N} \sum_{c=1}^{C} \mathrm{TP}_c. \tag{2}$$

*Weighted F1 score.* For each class $c$, precision, recall, and the F1 score are defined as

$$\mathrm{Precision}_c = \frac{\mathrm{TP}_c}{\mathrm{TP}_c + \mathrm{FP}_c}, \tag{3}$$

$$\mathrm{Recall}_c = \frac{\mathrm{TP}_c}{\mathrm{TP}_c + \mathrm{FN}_c}, \tag{4}$$

$$\mathrm{F1}_c = \frac{2 \cdot \mathrm{Precision}_c \cdot \mathrm{Recall}_c}{\mathrm{Precision}_c + \mathrm{Recall}_c}. \tag{5}$$

The weighted F1 score is computed as a support-weighted average across classes:

$$\mathrm{F1}_{\mathrm{weighted}} = \sum_{c=1}^{C} \frac{n_c}{N} \mathrm{F1}_c. \tag{6}$$

*Macro F1 score.* The Macro F1 score assigns equal weight to each class and is defined as

$$\mathrm{F1}_{\mathrm{macro}} = \frac{1}{C} \sum_{c=1}^{C} \mathrm{F1}_c. \tag{7}$$

*Overall classification score* To provide a single summary metric that jointly reflects overall accuracy, performance on rare cell types, and robustness to class imbalance, we define an overall classification score as:

$$S_c = Mean(\mathrm{Accuracy}, \mathrm{F1}_{\mathrm{macro}}, \mathrm{F1}_{\mathrm{weighted}}). \tag{8}$$

##### 1.4 Leiden subclustering and Fisher's exact test

For each annotated cell type, we performed Leiden subclustering in the latent embedding space to characterize intra-cell-type heterogeneity. Marker genes were identified for each Leiden subcluster using differential expression analysis within the corresponding cell type. To evaluate whether prototype distance captured transcriptional programmes associated with these subclusters, we compared prototype-distance-correlated genes with three marker-gene sets.

For a given cell type, let $D$ denote the set of prototype-distance-correlated genes, and let $U$ denote the background gene universe, defined as the set of genes tested in both the prototype-distance correlation analysis and the marker-gene analysis. We considered three marker-gene sets. First, let $M_{\mathrm{Leiden}}$ denote the union of marker genes from all Leiden subclusters within the cell type. The overlap proportion was calculated as

$$P_{\mathrm{Leiden}} = \frac{|D \cap M_{\mathrm{Leiden}}|}{|M_{\mathrm{Leiden}}|}. \tag{9}$$

Second, we identified the prototype-containing Leiden subcluster and denoted its marker-gene set as $M_{\mathrm{proto}}$. The overlap proportion between prototype-distance-correlated genes and marker genes of the prototype-containing subcluster was calculated as

$$P_{\mathrm{proto}} = \frac{|D \cap M_{\mathrm{proto}}|}{|M_{\mathrm{proto}}|}. \tag{10}$$

Third, for each non-prototype Leiden subcluster, marker genes were identified by comparing that subcluster against the prototype-containing subcluster. The union of these marker genes was denoted as $M_{\mathrm{nonproto-vs-proto}}$. The corresponding overlap proportion was calculated as

$$P_{\mathrm{nonproto-vs-proto}} = \frac{|D \cap M_{\mathrm{nonproto-vs-proto}}|}{|M_{\mathrm{nonproto-vs-proto}}|}. \tag{11}$$

To assess whether prototype-distance-correlated genes were significantly enriched in each marker-gene set, we performed a one-sided Fisher's exact test. For each marker-gene set $M$, where

$$M \in \{M_{\mathrm{Leiden}}, M_{\mathrm{proto}}, M_{\mathrm{nonproto-vs-proto}}\},$$

we constructed the following $2 \times 2$ contingency table:

| | Prototype-distance-correlated genes | Other genes |
|---|---|---|
| Marker genes in $M$ | $\lvert D \cap M\rvert$ | $\lvert M \setminus D\rvert$ |
| Non-marker genes | $\lvert D \setminus M\rvert$ | $\lvert U \setminus (D \cup M)\rvert$ |

The alternative hypothesis was that marker genes were enriched for prototype-distance-correlated genes:

$$H_1 : \text{odds ratio} > 1. \tag{12}$$

Therefore, Fisher's exact test was performed with a one-sided "greater" alternative. The resulting $P$-values were adjusted across cell types using the Benjamini–Hochberg procedure, and marker-gene sets with adjusted $q < 0.05$ were considered significantly enriched for prototype-distance-correlated genes.

#### 2 Supplementary Figures

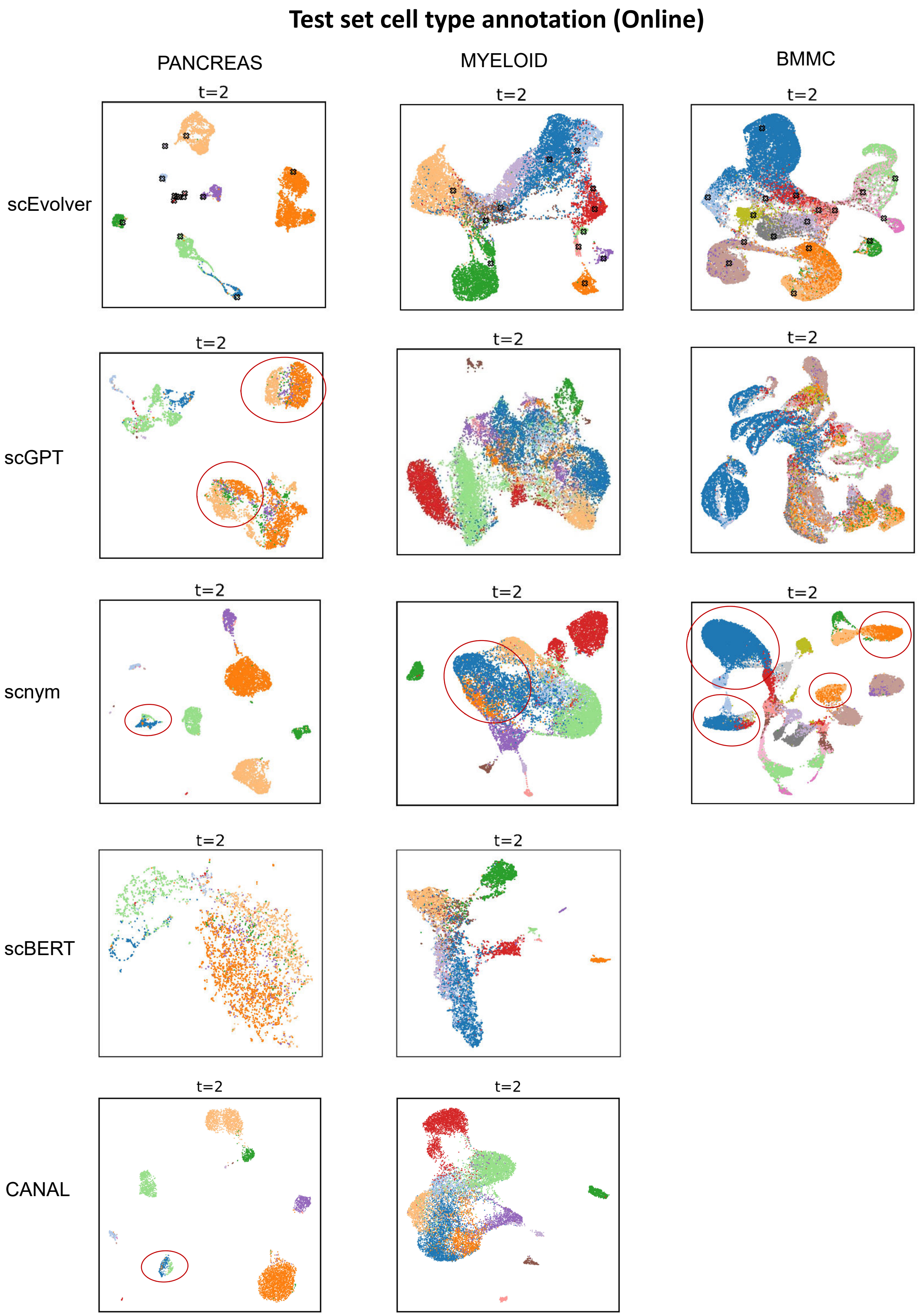


**Fig. S1 | Test set cell type annotation results(Online).** From left to right, panels show results for the PANCREAS, MYELOID, and BMMC datasets across different methods after continual learning. Cell-type annotations are evaluated on all test sets. scEvolver shows consistent alignment of cells with their biological labels, whereas other methods exhibit varying levels of batch effects. scBERT does not support multimodal data and is therefore not evaluated on the BMMC dataset.

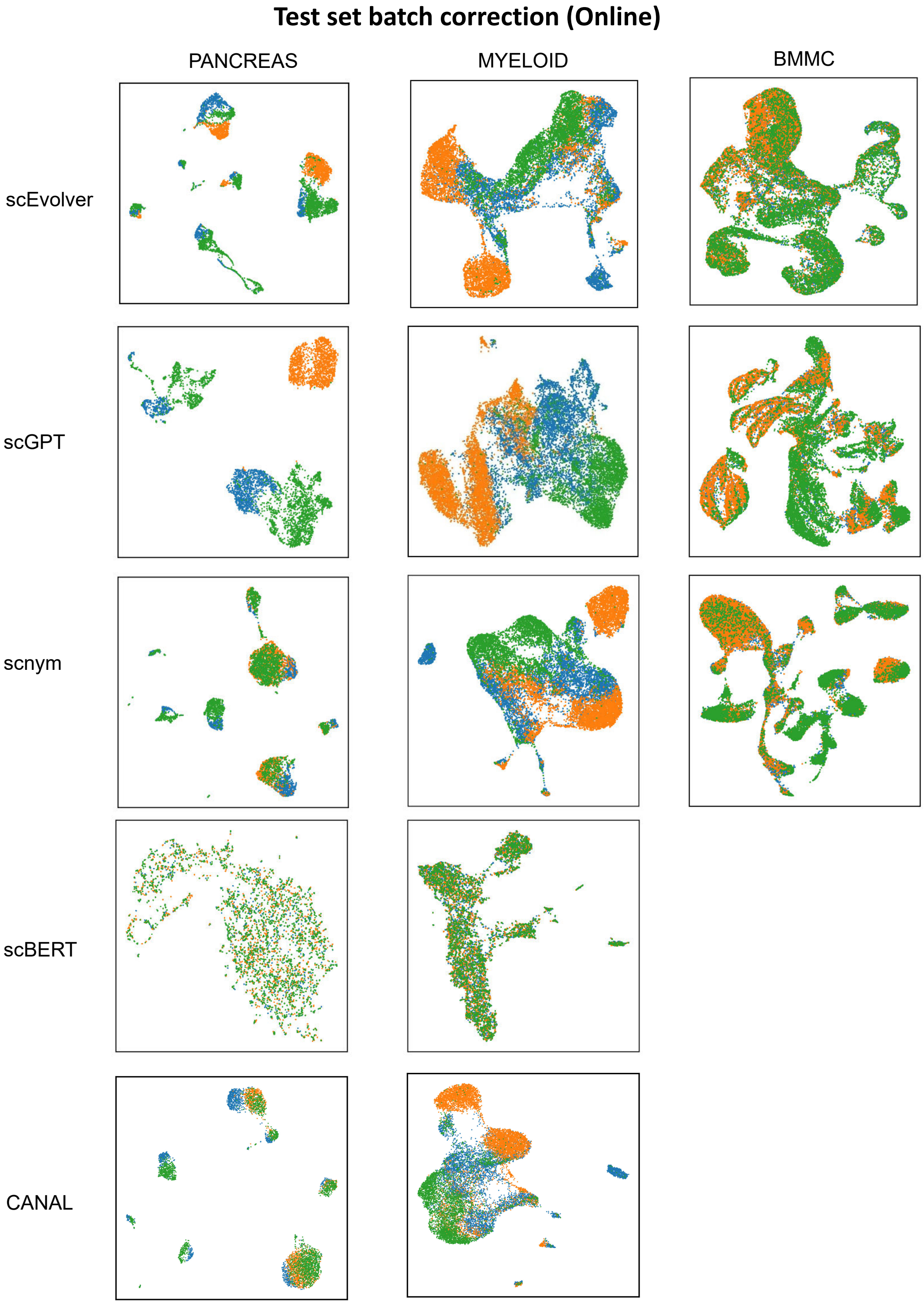


**Fig. S2 | Test-set batch correction results(Online).** From left to right, panels show results for the PANCREAS, MYELOID and BMMC datasets across different methods after continual learning. The latent embedding space is visualized to illustrate the degree of mixing across datasets, with greater mixing generally reflecting more effective batch effect mitigation. scEvolver shows reduced batch-associated separation compared with other approaches. scBERT does not support multimodal data and is therefore not evaluated on the BMMC dataset.

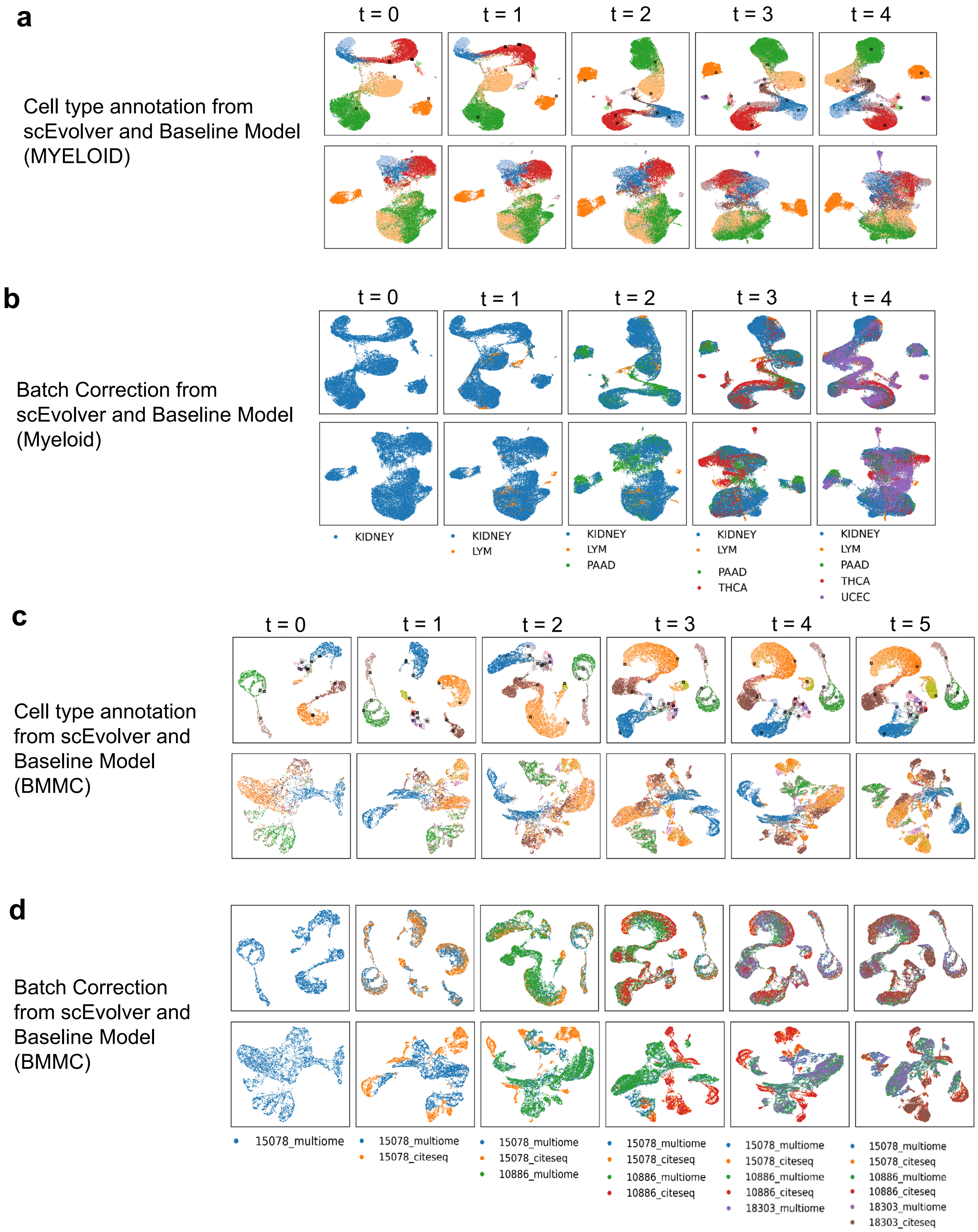


**Fig. S3 | scEvolver aligns representations across multi-tissues and multi-modalities, respectively. a, c,** UMAP visualization of cell embeddings learned by the continual learning model as data are progressively incorporated. Black crosses denote the locations of class prototypes in the latent space. **b, d,** UMAP visualization of batch mixing at different stages of online learning, with scEvolver showing improved batch mixing.

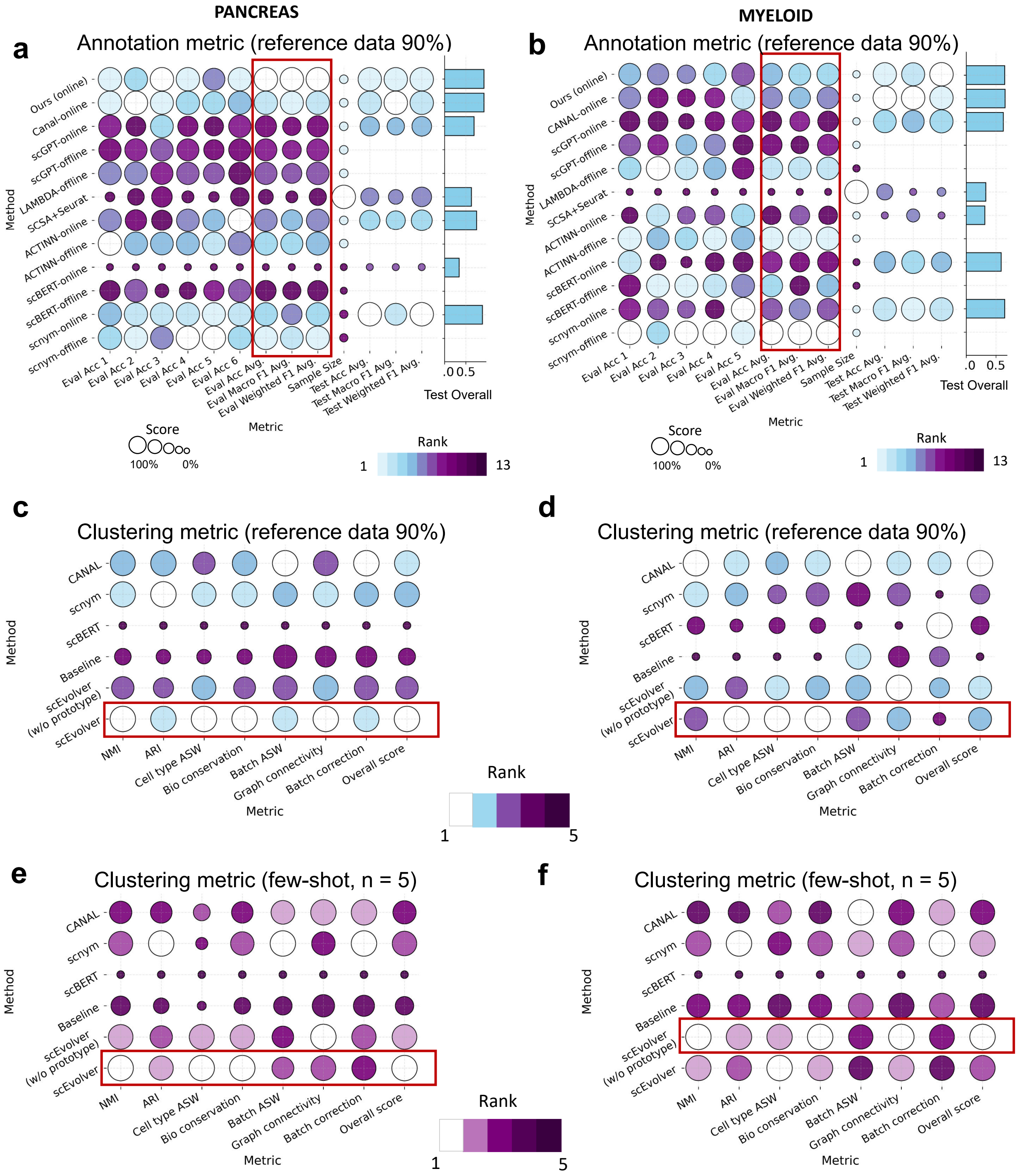


**Fig. S4 | Performance benchmark of reference and query test sets. a-b,** Annotation metrics on the evaluation and held-out query test sets after integrating all available reference training data (90% training data proportion). scEvolver consistently demonstrated strong generalization with minimal forgetting during continual learning. On the reference sets, scEvolver showed performance comparable to offline training, indicating effective retention of previously learned knowledge. In the online evaluation setting, scEvolver ranked first among all compared methods, highlighting its robustness under sequential data learning. **c-d,** Clustering metrics on the query test sets after integrating all available reference training data (90% training data proportion). **e-f,** Clustering metrics on the query test sets under a few-shot setting, using five labeled reference cells per cell type.

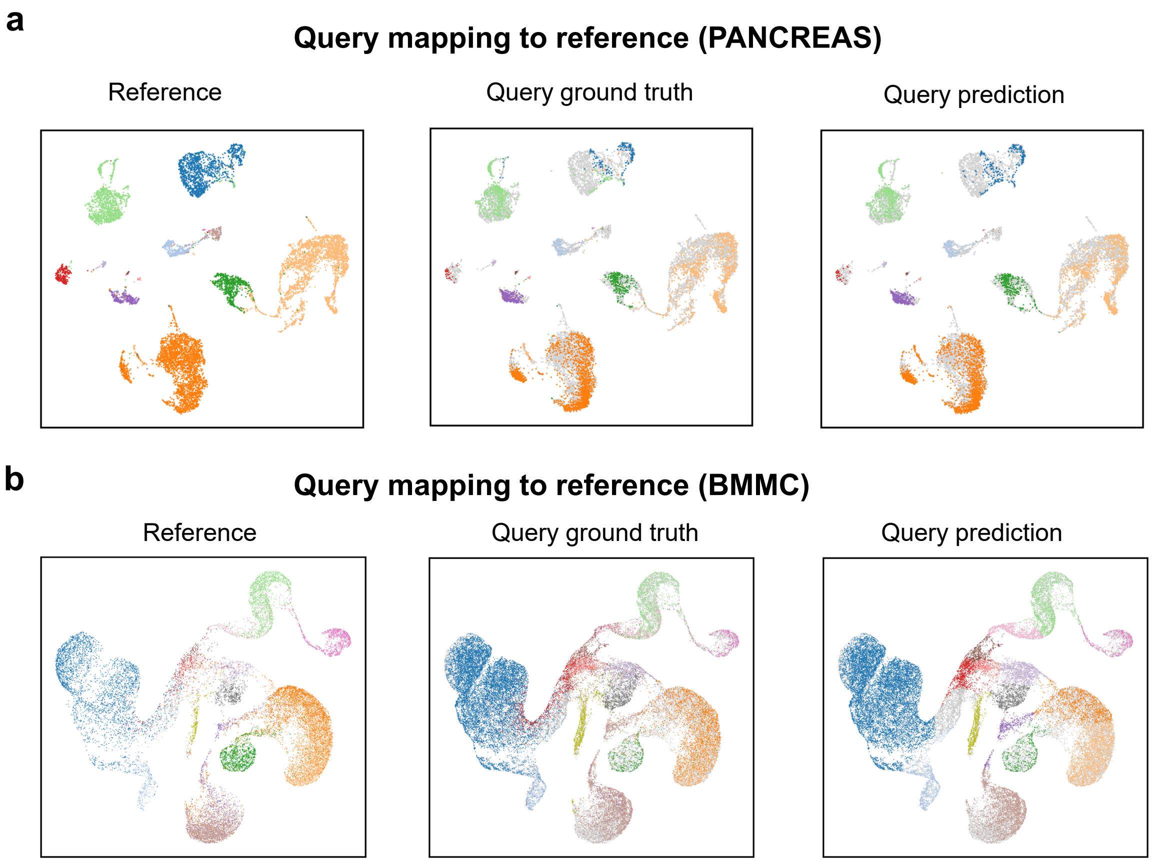


**Fig. S5 | Query mapping to reference in cross-platform and cross-modality datasets. a, b,** UMAP visualization of the reference data, query cells with ground-truth labels, and query cells with predicted annotations in the learned latent space, demonstrating accurate cell-type mapping by scEvolver.

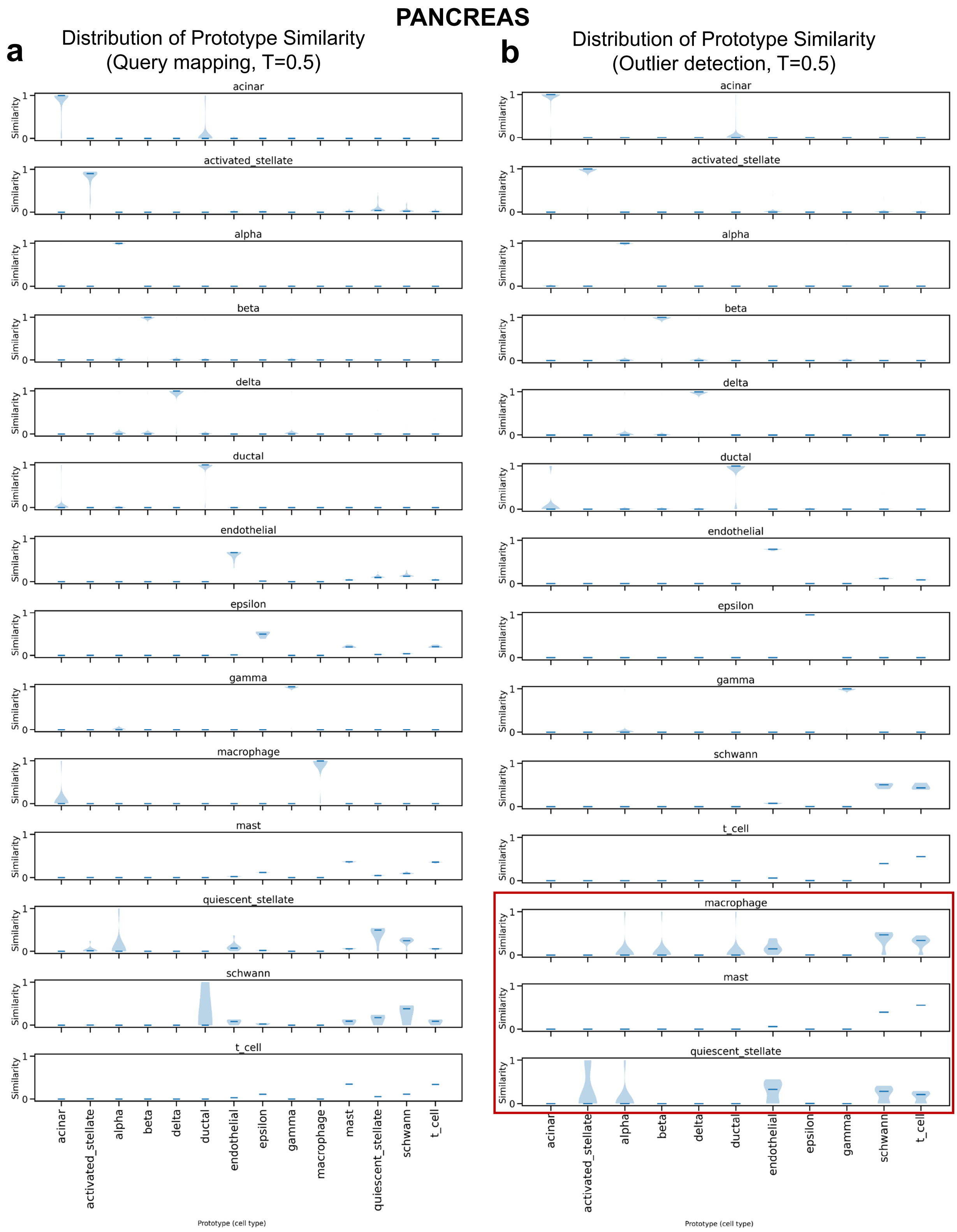


**Fig. S6 | Distribution of Prototype Similarity in PANCREAS dataset. a.** Query mapping results showing the distribution of similarity scores between query cells and their corresponding class prototypes. **b.** Outlier detection based on prototype similarity, where cells exhibiting low similarity to all reference prototypes are identified as potential out-of-distribution or unseen cell populations (highlighted by the red box).

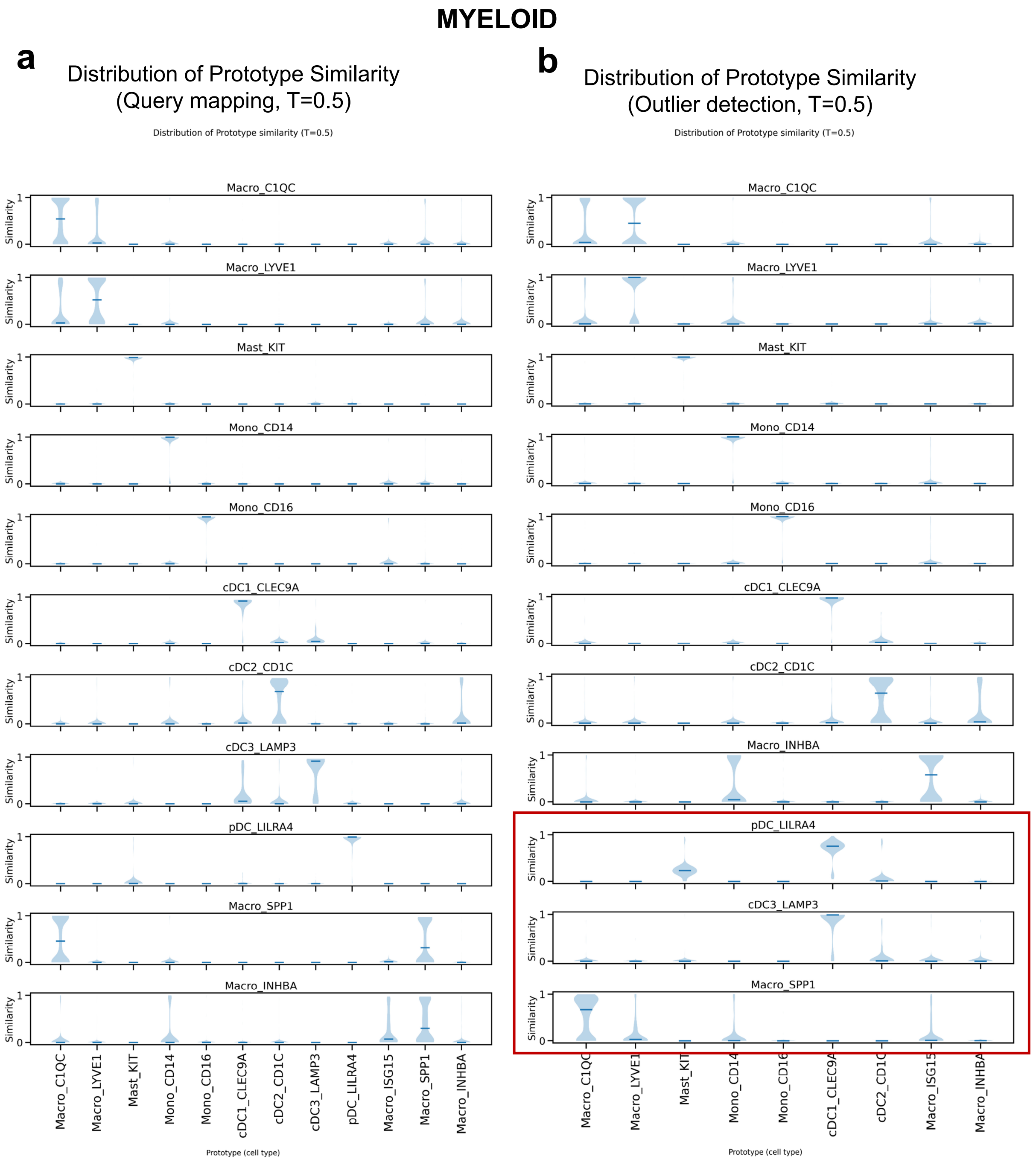


**Fig. S7 | Distribution of Prototype Similarity in MYELOID dataset. a.** Query mapping results showing the distribution of similarity scores between query cells and their corresponding class prototypes. **b.** Outlier detection based on prototype similarity, where cells exhibiting low similarity to all reference prototypes are identified as potential out-of-distribution or unseen cell populations (highlighted by the red box).

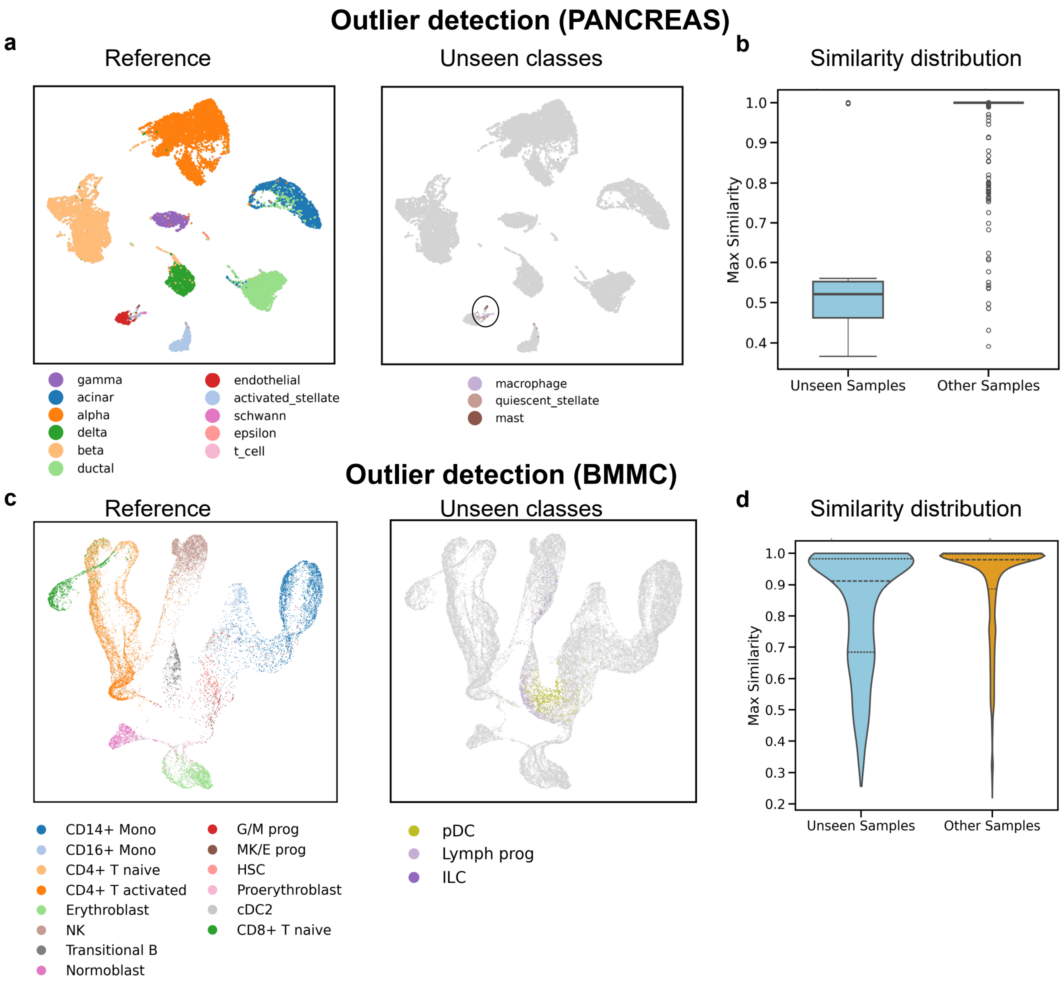


**Fig. S8 | Outlier detection in PANCREAS and BMMC datasets. a, c,** In the PANCREAS dataset, unseen cell types are clearly separated from the reference data in the UMAP embedding and exhibit distinct distributions of maximum prototype similarity compared with known classes. While in the BMMC dataset, unseen cell types map to regions distinct from reference clusters while remaining connected to the reference manifold, reflecting a continuous transition. Notably, clusters corresponding to HSCs, lymphoid progenitors, pDCs, and transitional B cells are positioned adjacently in the latent space, indicating continuity in the learned representations. This phenomenon is consistent with underlying transcriptional similarity and known lineage relationships. **b, d,** Distribution of maximum prototype similarity scores for known (Other Samples) and unseen cell types (Unseen Samples). Unseen cell types exhibit distinct distributions of maximum prototype similarity compared with known classes.

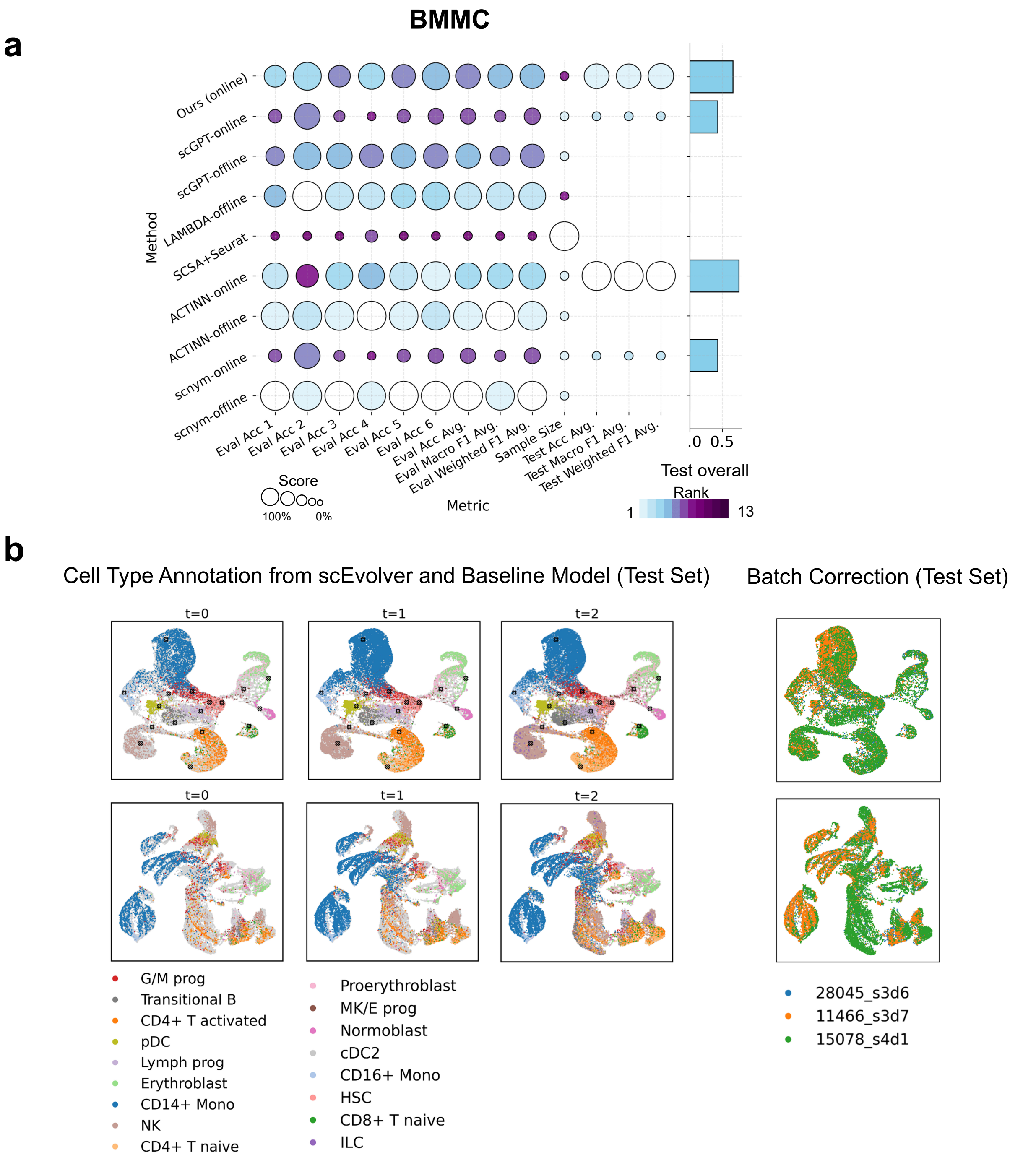


**Fig. S9 | Benchmark results on the BMMC datasets. a.** Performance evaluation on both reference and query test sets. scEvolver ranked second among all compared approaches, demonstrating robust performance under sequential data integration. Consistent with these quantitative results, UMAP visualizations of the test set show improved alignment and reduced batch effects compared with baseline (b) and other methods (supplementary Figs. S1 and S2).

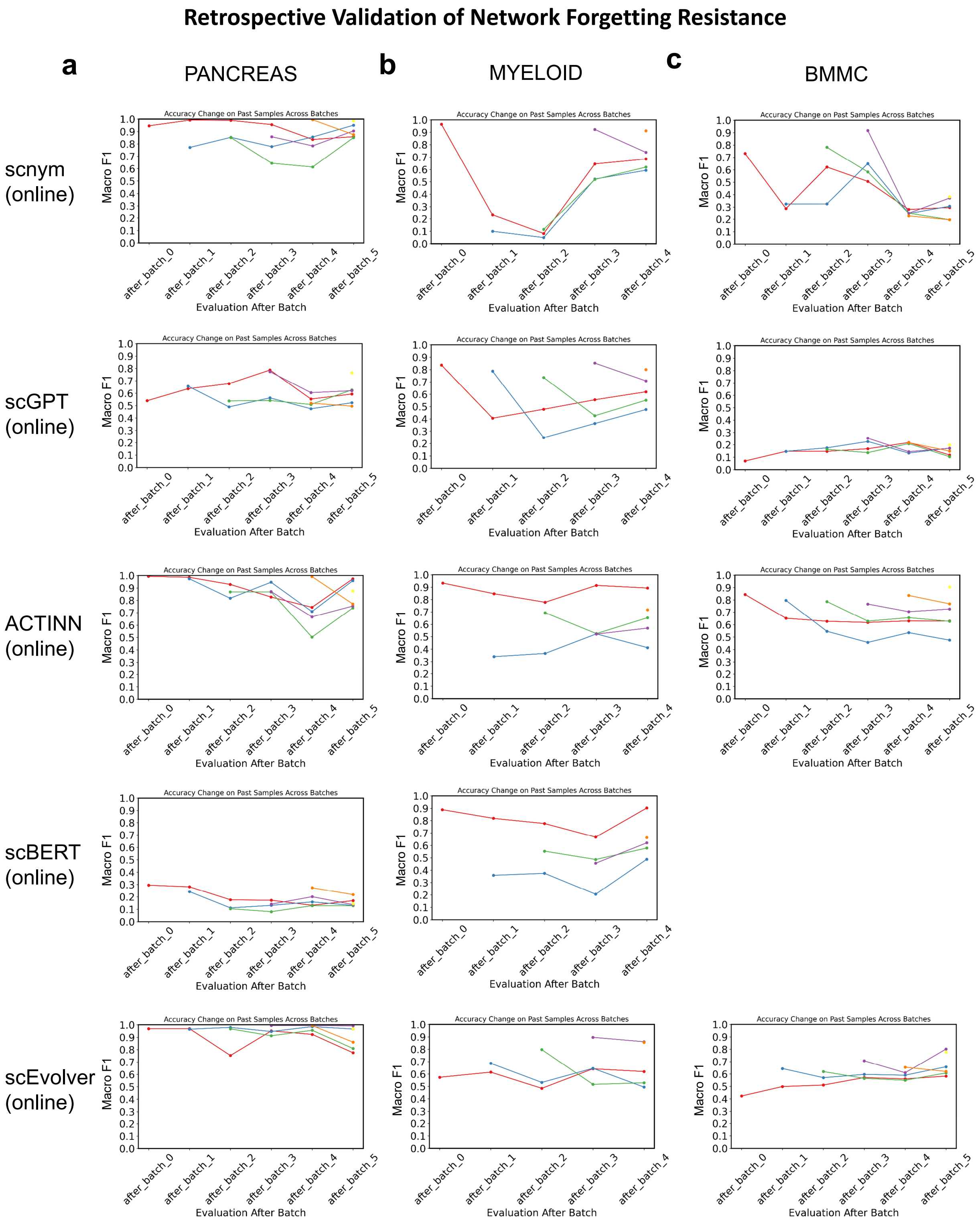


**Fig. S10 | Retrospective Validation of Network Forgetting Resistance.** Forgetting curves showing model macro-F1 on previously seen datasets during continual learning. While other methods exhibit varying degrees of forgetting across datasets, our method achieves the best balance between knowledge accumulation and forgetting resistance across all three scenarios. Each plot shows retrospective evaluation across learning stages, with the x-axis representing data accumulation.

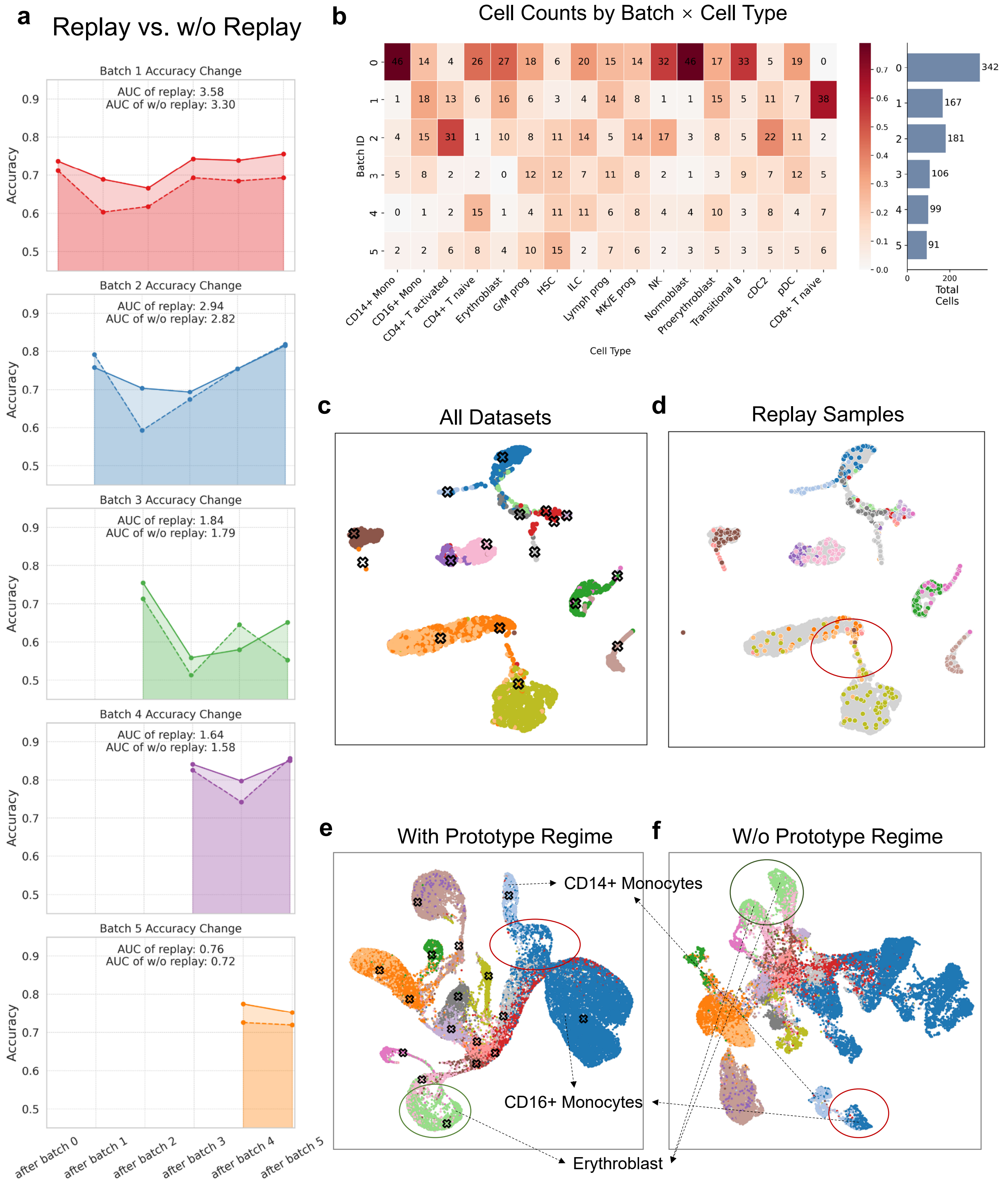


**Fig. S11 | Ablation analysis of memory replay and prototypes in scEvolver. a.** Contribution of memory replay to mitigating performance fluctuations caused by catastrophic forgetting and facilitating knowledge accumulation. Solid lines represent forgetting curves obtained with memory replay, whereas dashed lines show models trained without replay. **b.** Counts of replayed samples per cell type across batches. **c, d.** Compared to the distribution of all samples, replayed samples are more concentrated near class boundaries in the latent space. **e, f.** Latent-space clustering results with and without prototype guidance, showing that incorporating prototypes leads to smoother state transitions and improved continuity in the latent space.

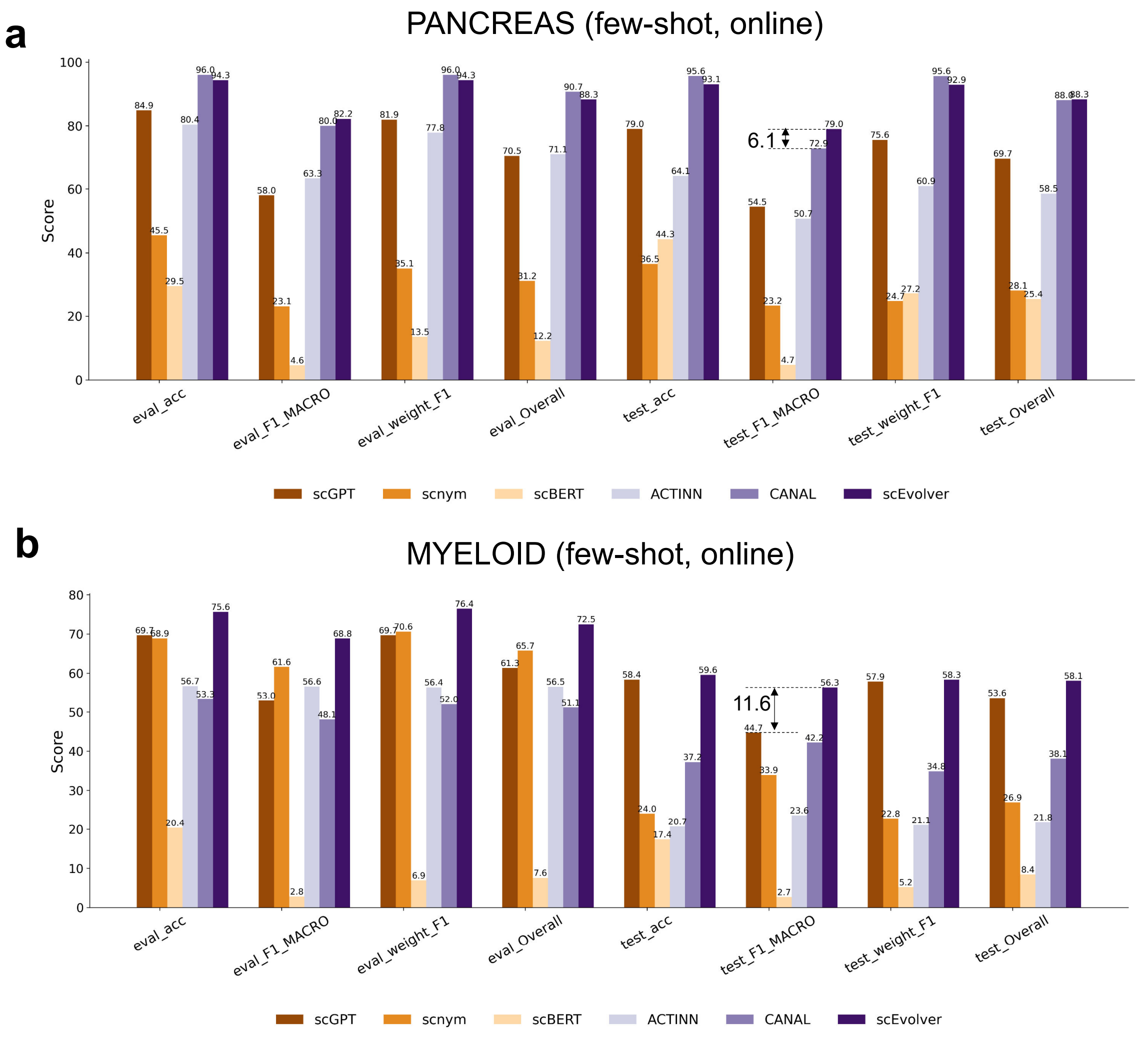


**Fig. S12 | Performance on evaluation sets and independent query test sets under a few-shot setting. a, b.** Evaluation on independent query test sets for the PANCREAS and MYELOID benchmarks, respectively, under the same few-shot regime. scEvolver consistently outperforms other approaches, including offline-trained models, achieving increases of 24.5% (PANCREAS) and 11.6% (MYELOID) in macro-F1 score, along with corresponding improvements of 20.3% and 6.2% in overall test score (see Methods).

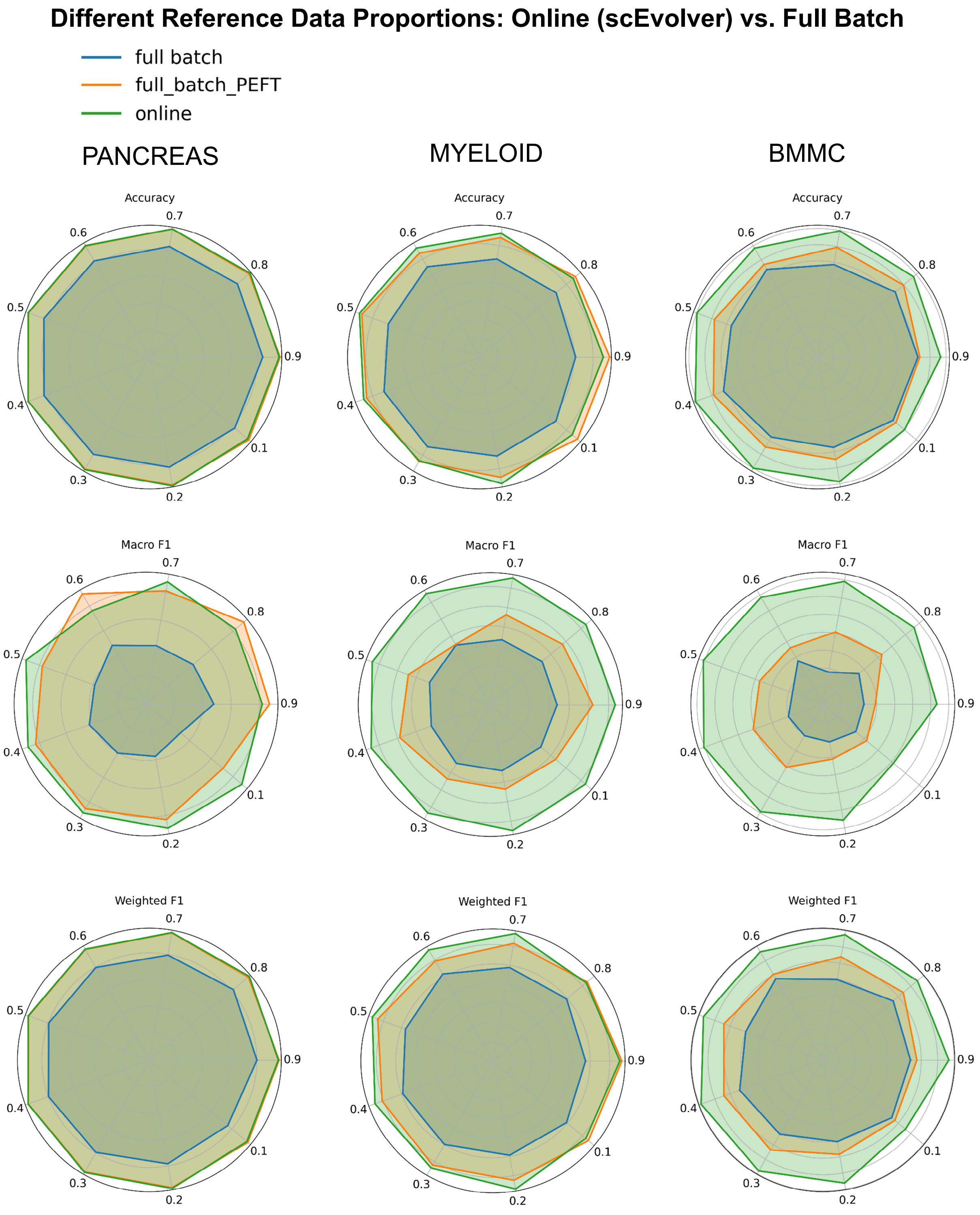


**Fig. S13 | Performance comparison with varying proportions of labeled reference data.** Comparison of scEvolver against full-batch fine-tuning (with and without PEFT) across three scenarios using 10–90% of labeled reference samples. scEvolver demonstrates stable performance and higher Macro F1 scores across all data proportions, indicating effective generalization with fewer labeled samples.

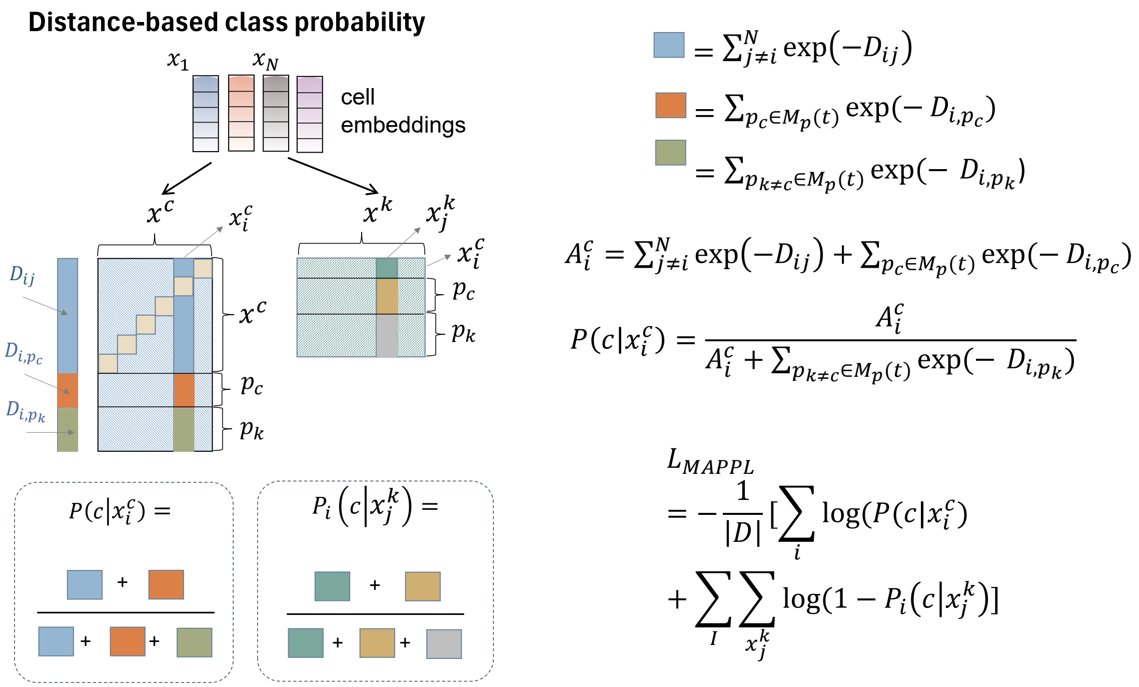


**Fig. S14 | A schematic illustration of distance-based class probability computation and the MAPPL loss.**

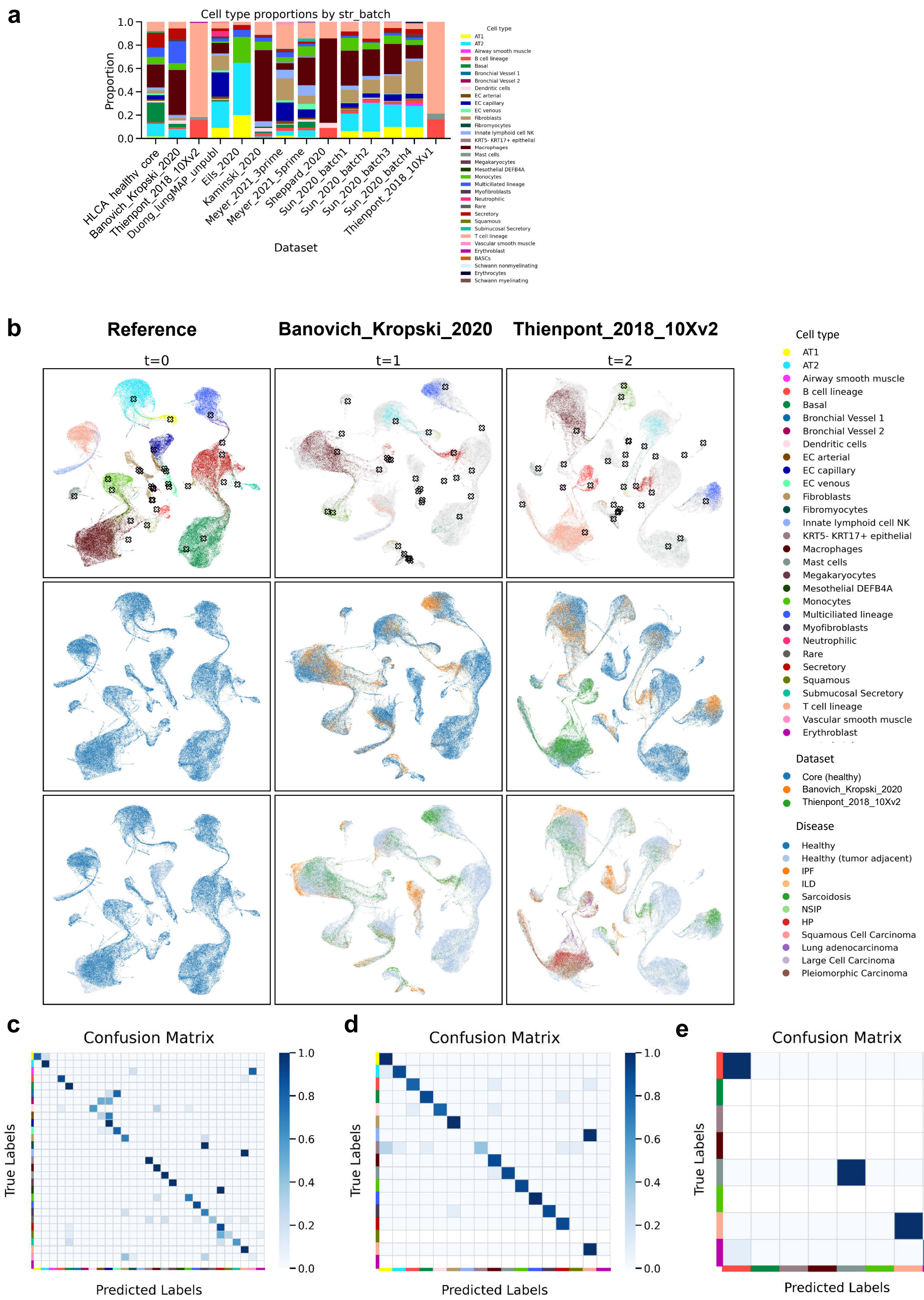


**Fig. S15 | Dataset composition and reference mapping of external lung datasets. a,** Cell-type composition across the HLCA reference and externally mapped lung datasets. Bar heights indicate the proportion of cells assigned to each annotated cell type within each dataset. **b,** UMAP visualization of the healthy reference and representative external datasets mapped into the prototype-anchored latent space. Cells are coloured by cell type, dataset identity and lung condition, showing that external datasets are projected into the shared reference space while retaining disease- and dataset-specific structure. **c–e,** Normalized confusion matrices showing label-transfer performance for the reference and mapped external datasets. Rows indicate true labels and columns indicate predicted labels; colour intensity denotes the fraction of cells assigned to each predicted class.

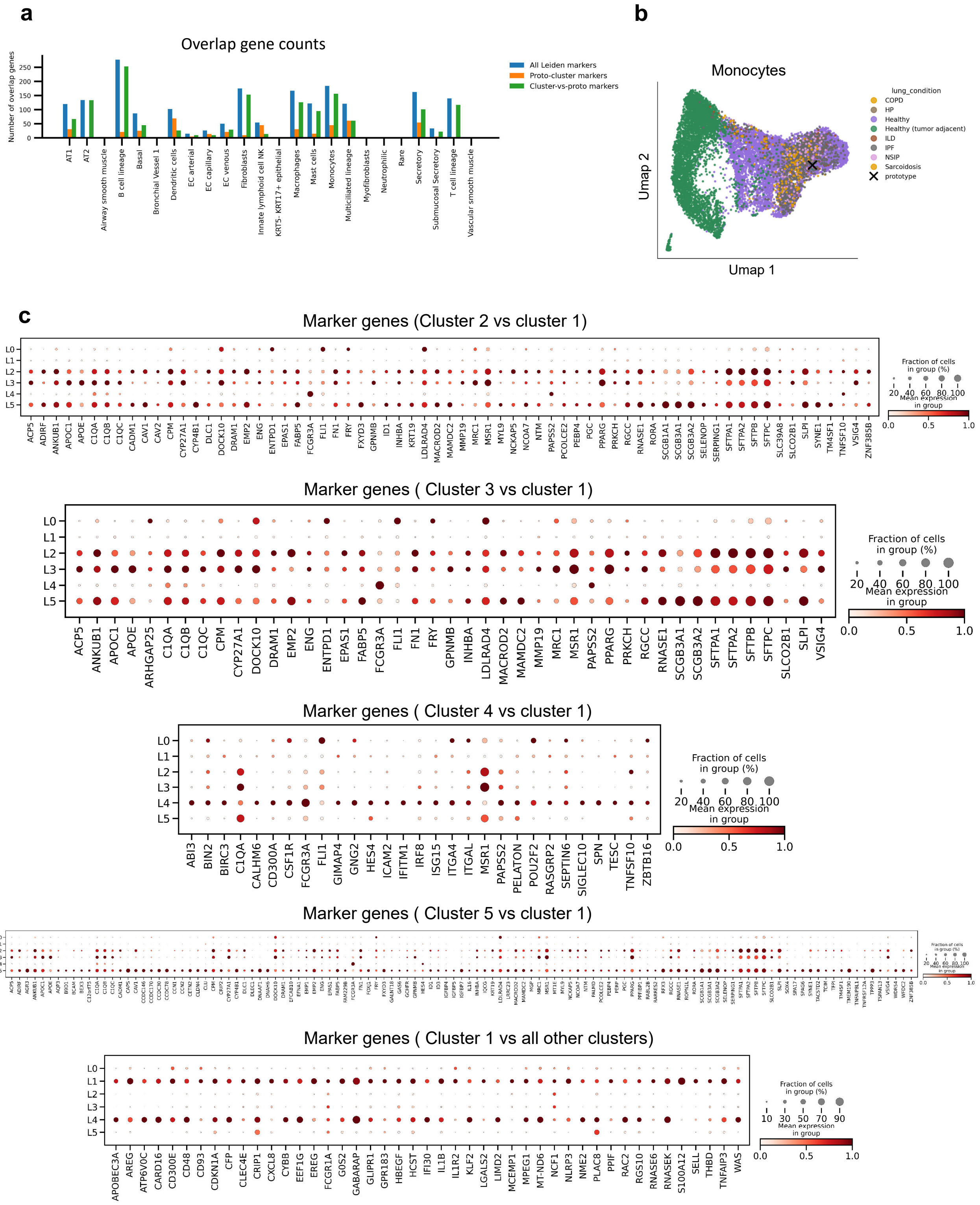


**Fig. S16 | Monocyte subcluster marker genes and prototype-distance-related transcriptional programmes. a,** Number of overlapping genes between prototype-distance-correlated genes and Leiden subcluster marker genes across cell types. Bars show overlaps with all Leiden markers, prototype-containing subcluster markers and markers distinguishing non-prototype subclusters from the prototype-containing subcluster. **b,** UMAP visualization of monocyte-labelled cells in the prototype-anchored latent space, coloured by lung condition. The reference monocyte prototype is indicated by the cross. **c,** Dot plots showing marker genes for monocyte Leiden subclusters. Marker genes were identified by comparing each non-prototype subcluster against the prototype-containing subcluster L1, and by comparing L1 against all other subclusters. Dot size indicates the fraction of cells expressing each gene, and colour indicates mean expression within each subcluster.
